# Convergence of multiple RNA-silencing pathways on GW182/TNRC6

**DOI:** 10.1101/2023.04.20.537632

**Authors:** Thomas Welte, Alison Goulois, Michael B. Stadler, Daniel Hess, Charlotte Soneson, Anca Neagu, Chiara Azzi, Marlena J. Lübke, Jan Seebacher, Isabel Schmidt, David Estoppey, Florian Nigsch, John Reece-Hoyes, Dominic Hoepfner, Helge Großhans

## Abstract

The RNA-binding protein TRIM71/LIN-41 is a phylogenetically conserved developmental regulator that functions in mammalian stem cell reprogramming, brain development and cancer. TRIM71 recognizes target mRNAs through hairpin motifs and silences them through molecular mechanisms that await identification. Here, we uncover that TRIM71 silences its targets by recruiting TNRC6/GW182, a core component of the miRNA-induced silencing complex (miRISC). Co-immunoprecipitation reveals interaction of TNRC6A with additional RNA-binding proteins and we demonstrate that AGO2, TRIM71, and UPF1 each recruit TNRC6 to specific, largely distinct sets of transcripts to silence them. As cellular TNRC6 levels are limiting, competition occurs among the silencing pathways, such that loss of AGO2 protein, or of AGO2 binding to TNRC6, enhances the activities of the other pathways. We conclude that a miRNA-like silencing activity is shared among different mRNA silencing pathways and that use of TNRC6 as a central hub provides a means to integrate their activities.

## Highlights

- A genome-wide CRISPR-screen reveals enhanced TRIM71-mediated RNA silencing activity upon miRNA pathway disruption
- The miRISC core component GW182/TNRC6 functions in AGO2-, TRIM71- and UPF1-mediated RNA silencing
- AGO2, TRIM71 and UPF1 recruit TNRC6A to specific sets of target transcripts
- AGO2 antagonizes TRIM71 and UPF1 activity by sequestering GW182/TNRC6

## Introduction

TRIM71/LIN-41 is a phylogenetically conserved RNA-binding protein (RBP) and E3 ubiquitin ligase that regulates stem and progenitor cell fates in diverse cell types and organisms (Chen et al., 2012; Ecsedi and Grosshans, 2013; Wang et al., 2019; Worringer et al., 2014). In mice, *Trim71* knock-out causes defects in mouse brain development, an exencephalus phenotype, defects in neural tube formation (Chen et al., 2012; Maller Schulman et al., 2008; Mitschka et al., 2015), germ cell loss (Torres-Fernández et al., 2021a), and death during mid-embryogenesis.

In humans, different heterozygous missense mutations in *TRIM71* have been linked to the etiology of congenital hydrocephalus (CH) due to premature differentiation of neuroprogenitor cells (Duy et al., 2022; Furey et al., 2018; Jin et al., 2020; Welte et al., 2019). All these mutations map to the NHL repeat domain through which TRIM71 recognizes trinucleotide-loop hairpins with specific sequence features (Kumari et al., 2018; Welte et al., 2019). When binding to mRNAs containing such hairpins in their 3’ untranslated regions (UTRs), TRIM71 induces transcript degradation, possibly accompanied by some level of translational repression (Chang et al., 2012; Loedige et al., 2013; Mitschka et al., 2015; Welte et al., 2019).

The molecular machinery that mediates TRIM71 silencing is largely unknown. In mouse embryonic stem cells (mESCs), TRIM71 was shown to cooperate with miRNAs to achieve repression of the *Cdkn1a* mRNA (Chang et al., 2012). However, miRNAs do not appear to be generally required for TRIM71 activity (Mitschka et al., 2015; Torres-Fernández et al., 2019; Welte et al., 2019). Indeed, silencing of *CDKN1A* mRNA by TRIM71 is conserved in human HEK293T and HepG2 cells, but does not rely on an intact miRNA pathway (Torres-Fernández et al., 2019). Instead, the RNA binding proteins UPF1 and SMG1 were reported to promote silencing of *CDKN1A* and possibly other TRIM71 targets in these cells (Torres-Fernández et al., 2019). The molecular basis of this apparent cooperation, and the extent to which it is required for TRIM71 activity, have remained unknown.

Using a CRISPR-based screen to uncover the molecular mode of action of TRIM71, we find that TNRC6/GW182 is a key mediator of its activity. In mice and humans, TNRC6 comprises a set of three paralogous proteins that function redundantly in the miRNA pathway as a core component of the miRNA silencing complex (miRISC) (Ding and Han, 2007; Eulalio et al., 2008; Jakymiw et al., 2005; Liu et al., 2005). (Throughout this paper, we will generically refer to TNRC6 unless we manipulate experimentally a specific isoform.) Binding to an Argonaute protein (AGO1∼4) associated with a miRNA facilitates recruitment of TNRC6 to target transcripts in a sequence-specific manner. Through its interactions with a diversity of general RNA decay and silencing effector proteins, including the CCR4/NOT deadenylase complex, the PAN2/PAN3 deadenylase complex, cytoplasmic poly(A)-binding protein (PABPC) and the RNA helicase DDX6, TNRC6-binding then induces target transcript silencing through degradation or translational repression (Braun et al., 2011; Braun et al., 2013; Chekulaeva et al., 2011; Mathys et al., 2014).

We find that TRIM71 recruits TNRC6 to a largely distinct set of transcripts, not targeted by miRISC, to induce their silencing. We observe that TNRC6 levels are limiting for TRIM71 activity. Accordingly, uncoupling TNRC6 from AGO enhances TRIM71 activity. This miRISC-independent activity of TNRC6 extends to at least one other RBP, UPF1, identifying TNRC6 as a central hub of post-transcriptional gene regulation. Thus, it can integrate the activities of multiple silencing pathways, exemplified by miRISC’s ability to attenuate other RNA silencing pathways through sequestration of TNRC6.

## Results

### A genome-wide CRISPR/Cas9 sgRNA screen identifies potential co-factors and modulators of TRIM71 activity

To identify genes relevant for TRIM71-mediated gene regulation, we performed a genome-wide CRISPR/Cas9 sgRNA screen. Based on our previous studies on TRIM71 RNA target recognition (Welte et al., 2019), we constructed a minimal GFP reporter system in mESCs that recapitulates TRIM71-dependent gene regulation. To reduce the likelihood of our reporter being targeted by other endogenous regulatory circuits, we chose to construct a heterologous system based on *C. elegans* 3’UTRs. Using Cas9-expressing mESCs (**Supplemental Figure S1A-C**), we generated the following cell lines: *GFP_Ctrl* cells (negative control): The *C. elegans unc-54* 3’ UTR was fused to an *EGFP*-coding sequence. This UTR has no TRIM71-binding motifs and is thus not repressed by TRIM71, resulting in high GFP levels (**Figure 1A**).

**Figure 1.**
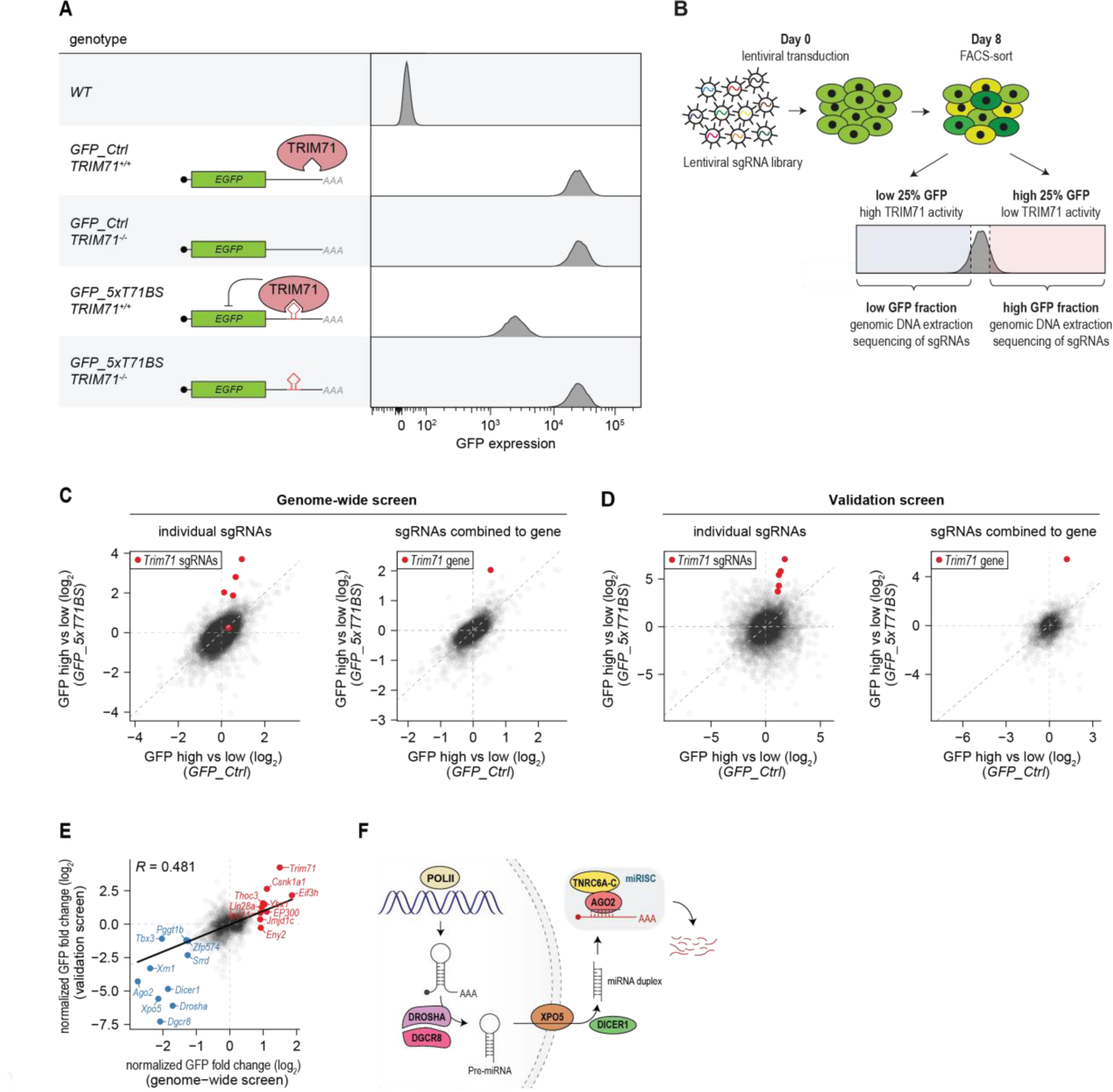
Two CRISPR/Cas9 sgRNA-screens identify genes involved in miRNA processing and silencing as potent repressors of TRIM71-dependent gene silencing. **(A)** Reporter cell lines used in this study (*left*, schematic depiction) and their GFP levels quantified by flow cytometry (*right)*. **(B)** Schematic depiction of the CRISPR/sgRNA screen. **(C), (D)** Scatter plots showing results of a genome-wide **(C)** and a validation CRISPR/sgRNA screen **(D)**, respectively. Enrichment of sgRNAs (left panels) in high vs low GFP fractions are shown for *GFP_Ctrl* cells (x-axis), and *GFP_5×T71BS* (y-axis) cells. *Trim71* sgRNAs are colored in red. The right panels show the aggregated (median) effect of all sgRNAs per gene. *Trim71* is colored in red. **(E)** Scatter plot showing a comparison of effects of genome-wide (x-axis) and validation sgRNA/CRISPR screens (y-axis) on gene-level. Effects of high vs. low GFP fractions in *GFP_5×T71BS* cells are normalized to the same effects in *GFP_Ctrl* cells (from **[C]** and **[D]**). The ten genes with the highest (marked in red) and the lowest (marked in blue) enrichment in the genome-wide CRISPR/sgRNA screen are indicated. A linear regression model was fitted to the data (shown as black line). *R* indicates the Pearson’s correlation coefficient. Data source for **(C-E)**: **Supplemental Table S1. (F)** Schematic drawing illustrating the key components of the miRNA pathway.

*GFP_5×T71BS* cells (positive control): A fragment of the *C. elegans mab-10* 3’UTR containing five hairpin motifs recognized by TRIM71(Kumari et al., 2018; Welte et al., 2019) was cloned into the *GFP::unc-54* 3’ UTR construct, leading to a marked reduction in GFP levels relative to *GFP_Ctrl* cells (**Figure 1A**). This repression depended on TRIM71, as disruption of *Trim71* in *GFP_5×T71BS* cells caused a shift to high GFP levels, equal to those observed in *GFP_Ctrl* cells. By contrast, *Trim71* disruption in *GFP_Ctrl* cells did not affect the signal (**Figure 1A, Supplemental Figure S1D).**

We used the *GFP_5×T71BS* cell line to perform a CRISPR/sgRNA screen(Hoepfner et al., 2019), seeking to identify conditions that would either impair TRIM71 activity, and thus increase GFP signal; or enhance TRIM71 activity, and thus decrease GFP signal. The *GFP_Ctrl* cell line allowed us to control for non-specific effects, not related to modulating TRIM71 activity.

Each cell line was separately transduced with two lentiviral libraries comprising in total 91,000 sgRNAs (∼5 sgRNAs/gene) at a multiplicity of infection (MOI) of 0.31 and ∼700-fold coverage per sgRNA. At day 8, cells were harvested and sorted by fluorescence-activated cell sorting (FACS) to fractions with low and high GFP levels, respectively (‘25% low’ and ‘25% high’; **Figure 1B**): DNA from each fraction, as well as an unsorted baseline control was extracted, followed by sequencing of sgRNAs. We aggregated sgRNAs by gene and quantified enrichment in the ‘25% high’ or ‘25% low’ fractions, respectively. We corrected for non-specific effects on gene expression subtracting enrichment values obtained for *GFP_Ctrl* cells from those obtained for *GFP_5×T71BS* cells (see **STAR Methods** for details).

Using the top and bottom 985 ranking genes, and 400 unchanged control genes, we performed a validation screen using a lentiviral library with 12,000 new sgRNAs (from (Michlits et al., 2020); **Supplemental Table S1**), transfected at an MOI of 0.1 and ∼500-fold coverage. Effects from both screens correlated well (*R* = 0.481). TRIM71 ranked second and first, respectively, among the ‘25% high’ genes in the two screens, validating the approach (**Figure 1C, D**).

Additional genes were reproducibly enriched in the ‘25% high’ and ‘25% low’ set. Strikingly, for the ‘25% low’ set, the top 5 genes were components of the miRNA pathway(Lin and Gregory, 2015), namely *Drosha* and *Dgcr8*, which together form the microprocessor complex that processes pri-miRNAs to pre-miRNAs; *Xpo5*, which mediates export of pre-miRNAs from the nucleus to the cytoplasm; *Dicer1*, which processes pre-miRNAs to generate miRNA duplexes; and *Ago2*, the core miRISC component that binds mature miRNAs (**Figure 1E, F**). This evident pathway enrichment suggested that our screen was close to saturation. It also pointed to a specific connection between the miRNA pathway and TRIM71, which we sought to explore further.

### The miRNA pathway attenuates TRIM71-dependent gene silencing independent of TRIM71 levels

We validated the effects seen in the screen through sgRNA enrichment by direct assessment of TRIM71 activity, using both the *GFP_5×T71BS* reporter and luciferase-based reporters containing either the *same* 5×T71BS 3’UTR or the 3’UTR of *Plxnb2*, a previously identified TRIM71 target (Mitschka et al., 2015; Welte et al., 2019). Knock-out of *Dicer1*, *Xpo5* or *Ago2* enhanced repression of each of the TRIM71 target reporters without altering expression of control reporters that were not regulated by TRIM71 (**Figure 2A-C, Supplemental Figure S2A**). Conversely, transfection of exogenous *Ago2* reverted *GFP_5×T71BS* reporter repression to the wild-type situation (**Figure 2D, Supplemental Figure S2B**). The effects were specific to the TRIM71 pathway, as changes in reporter repression only occurred in the presence of *Trim71*. We conclude that attenuation of TRIM71 activity through the miRNA pathway is reproducible and specific, and not limited to the heterologous reporter or GFP signal as a read-out.

**Figure 2.**
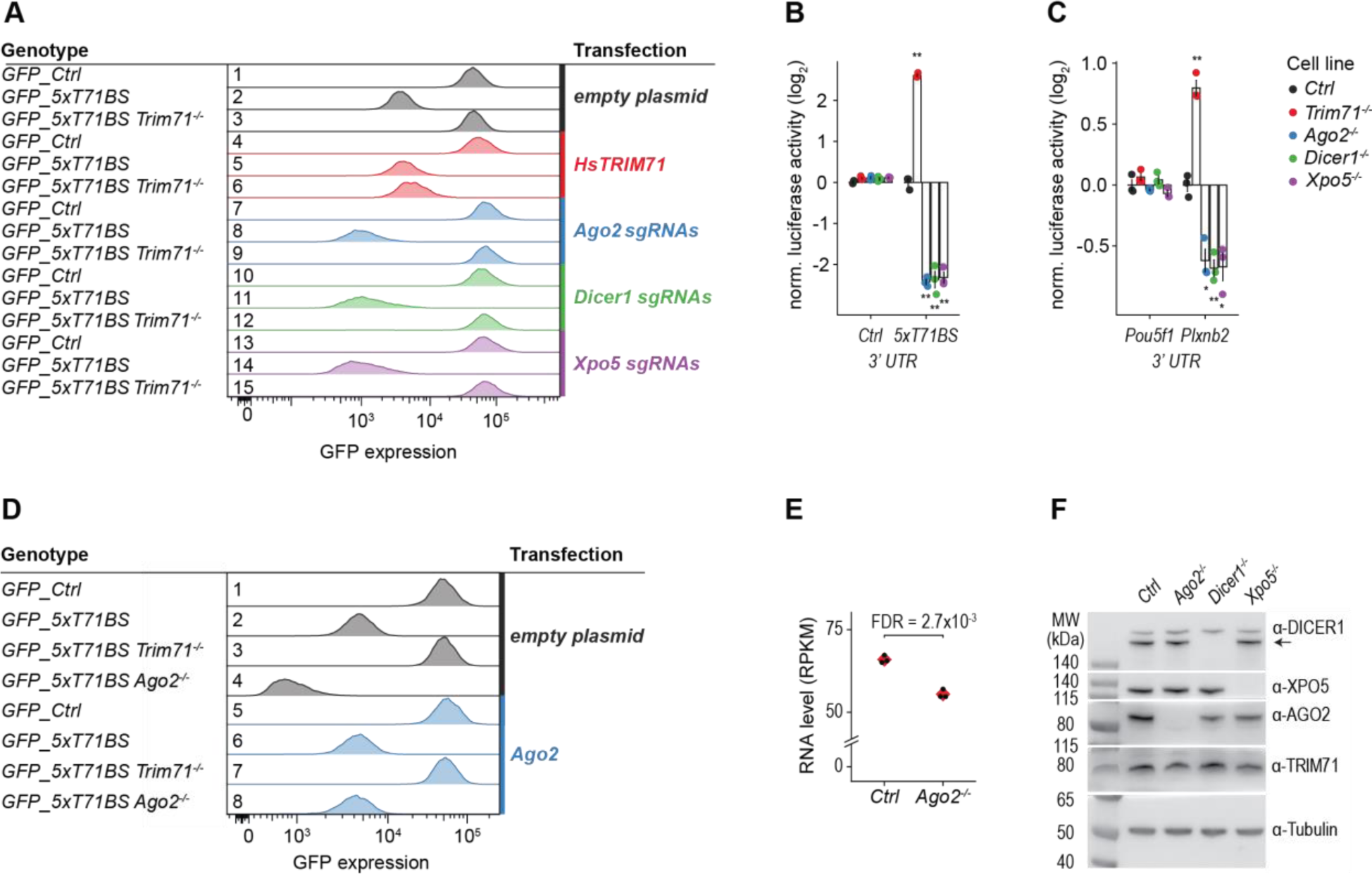
miRNA pathway genes act as repressors of TRIM71-dependent gene silencing. **(A, D)** Flow cytometry analysis showing the effect of sgRNAs targeting *Xpo5*, *Dicer1*, *Ago2*, and of transfection with *HsTRIM71* overexpression plasmids as indicated. **(B, C)** Luciferase reporter gene assay with the indicated *Renilla* 3′ UTR-fusion constructs in *WT*, *Trim71*, *Ago2*, *Dicer1*, and *Xpo5* knock-out mESCs. Data were normalized to *WT* and represent mean ± standard error of the mean (SEM). *n* = 3 biological replicates. Significance was tested using a two-tailed Student’s *t*-test. (*) *P* < 0.05; (**) *P* < 0.01; (***) *P* < 0.001; otherwise not significant (*P* ≥ 0.05). **(E)** Plot showing levels of *Trim71* mRNA in control, and *Ago2^-/-^* cells (*n* = 3 biological replicates; background cell line: *GFP_5×T71BS*). RNA levels are shown as reads per kilobase of transcript per million of mapped reads (RPKM). FDR: Benjamini Hochberg adjusted *P*-value. **(F)** Western blots showing the effect of sgRNAs targeting *Xpo5*, *Dicer1*, *Ago2* as indicated. Parental cell line (‘*Ctrl’*): *GFP_5×T71BS*.

LIN-41/TRIM71 has been shown to be repressed during development of *C. elegans* and mouse by the *let-7* family of miRNAs (Aeschimann et al., 2019; Ecsedi et al., 2015; Rybak et al., 2008; Slack et al., 2000). Although this repression normally occurs in differentiating cells, we wondered if the observed change of TRIM71 silencing activity simply reflected its release from miRNA-mediated repression. This was not the case: RNA sequencing revealed that *Trim71* mRNA levels did not increase in *Ago2^-/-^* relative to wild-type mESCs (**Figure 2E**), and we failed to observe an increase in TRIM71 protein levels upon knock-out of *Ago2*, *Dicer1*, and *Xpo5* when tested in two independent experiments (**Figure 2F, Supplemental Figure S2A**). By contrast, and as reported previously (Martinez and Gregory, 2013; Wang et al., 2020), Ago2-levels were reduced in *Dicer1*, and *Xpo5* knock-out cells (**Figure 2F, Supplemental Figure S2A**), consistent with the notion that Ago2 requires miRNA binding for stabilization (Smibert et al., 2013). Finally, overexpression of *Homo sapiens TRIM71* (**Supplemental Figure S2A**) complemented *Trim71* knock-out cells to restore silencing of the *GFP_5×T71BS* reporter (**Figure 2A**, row 3 vs. row 6), but did not further increase silencing of the reporter in *Trim71* wild-type cells (**Figure 2A**, row 2 vs. row 5).

We conclude that TRIM71 is not detectably repressed by miRNAs in mESCs, and that its levels are not limiting for its target silencing activity. Put differently, the modulation of TRIM71 activity through miRNA pathway disruption appears to occur downstream of TRIM71 protein accumulation.

### TNRC6 contributes to TRIM71-dependent gene silencing

Our sgRNA screens had identified the major components of miRNA biogenesis and AGO2, a core miRISC component (**Figure 1F**) and the most prominently expressed paralog of the four mammalian *Ago* genes(Boroviak et al., 2018; Lykke-Andersen et al., 2008). However, the second miRISC core component, TNRC6, was not identified in our sgRNA screens.

We suspected that TNRC6 was not identified because cells contain three TNRC6 isoforms, TNRC6A∼C, that are thought to function redundantly (Baillat and Shiekhattar, 2009; Johnson et al., 2021; Liu et al., 2019; Meister et al., 2005). Hence, we sought to directly test a possible function of TNRC6 in TRIM71-mediated RNA silencing. Despite extensive attempts, we failed to obtain *Tnrc6* triple-knockout cells, and no such cell line has been described in the literature, suggesting that TNRC6 is essential for mESC survival. By contrast, the miRNA pathway is dispensable for mESC survival (see results above), which implies that TNRC6 may have additional functions beyond their established role in the miRNA pathway.

To assess the consequences of TNRC6 deregulation for the TRIM71 pathway, we overexpressed *HsTNRC6A*, *B*, and *C* in the GFP reporter cell lines. This decreased GFP levels in *GFP_5×T71BS* cells (**Figure 3A**, row 5 vs rows 7-9). The effect was TRIM71-dependent, as no silencing was observed in *Trim71^-/-^* cells overexpressing *TNRC6* paralogs (**Figure 3A**). No changes in TRIM71 or AGO2 levels were observed upon *TNRC6* overexpression (**Figure 3B**). Thus, *TNRC6* overexpression is sufficient to recapitulate *Ago2* knock-out phenotypes and enhance TRIM71 activity.

**Figure 3.**
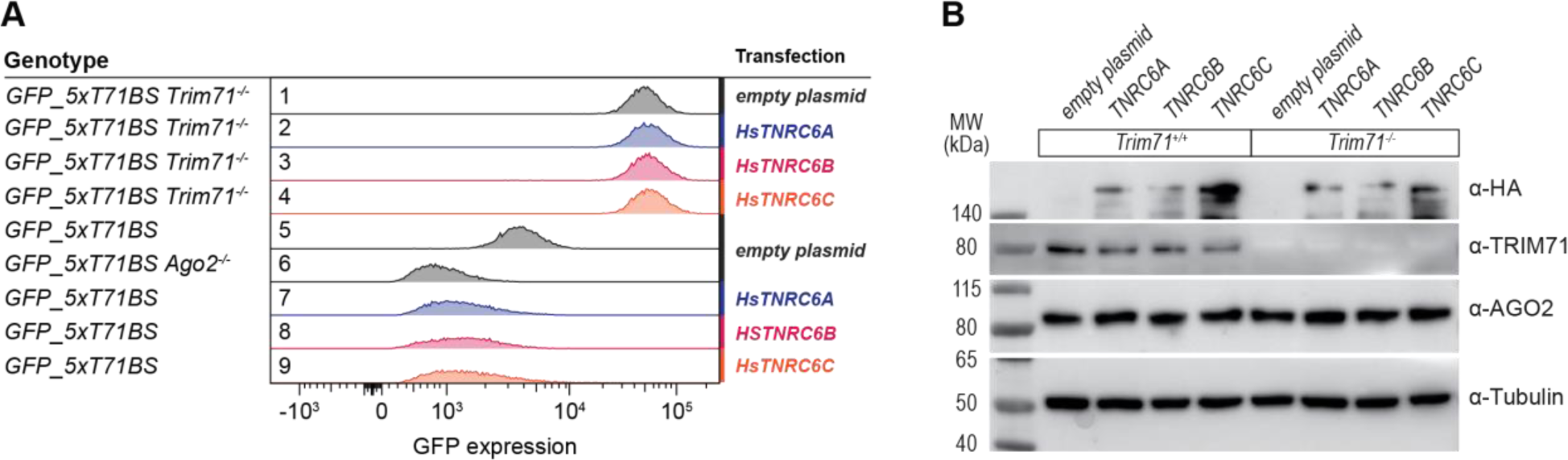
TNRC6 contributes to TRIM71-dependent gene silencing. **(A)** Flow cytometry analysis showing the effect of overexpressing *HsTNRC6A*, *B*, and *C* in *Trim71^+/+^* and *Trim71^-/-^ GFP_5×T71BS* cells, respectively. *GFP_5×T71BS Ago2^-/-^* (row 6) is included for comparison and replotted from above (Figure 2D, row 4). Transfected cells were selected with Puromycin for two days. **(B)** Western blots showing the expression of HA-tagged HsTNRC6A, B, and C (upper panel), and corresponding TRIM71, AGO2, and Tubulin levels of cells used in **(A)**.

We emphasize that the role of TNRC6 is opposite to that of AGO2 and the other miRNA pathway components in these assays, in that increased, rather than decreased TNRC6 levels enhance TRIM71 activity. Combined with our inability to observe protein level regulation of TRIM71 by the miRNA pathway, and the inability of *Trim71* overexpression to enhance TRIM71 activity, these data support a model where TNRC6 functions directly in TRIM71-mediated transcript repression. As loss of miRNAs both destabilizes AGO and reduces its affinity to TNRC6n (ref. (Elkayam et al., 2017)), a direct function of TNRC6 in the TRIM71 pathway could also explain why inactivation of miRNA biogenesis enhances TRIM71 activity.

### TNRC6 proteins coimmunoprecipitate with both AGO2 and TRIM71

We wondered whether the functional connection between TRIM71 and TNRC6 proteins reflected physical interactions. To test this possibility, we immunoprecipitated endogenously FLAG-AVI-tagged TRIM71 (ref. (Welte et al., 2019)) or endogenously FLAG-tagged AGO2 (ref. (Li et al., 2020)) mESCs. For both proteins, we consistently observed co-precipitation of TNRC6A and TNRC6B. Conversely, when we immunoprecipitated HA-AVI-tagged endogenous TNRC6A, we observed co-precipitation of both AGO2 and TRIM71 (**Figure 4A-C**). Hence, TRIM71 and AGO2 both interact biochemically with TNRC6A/B.

**Figure 4.**
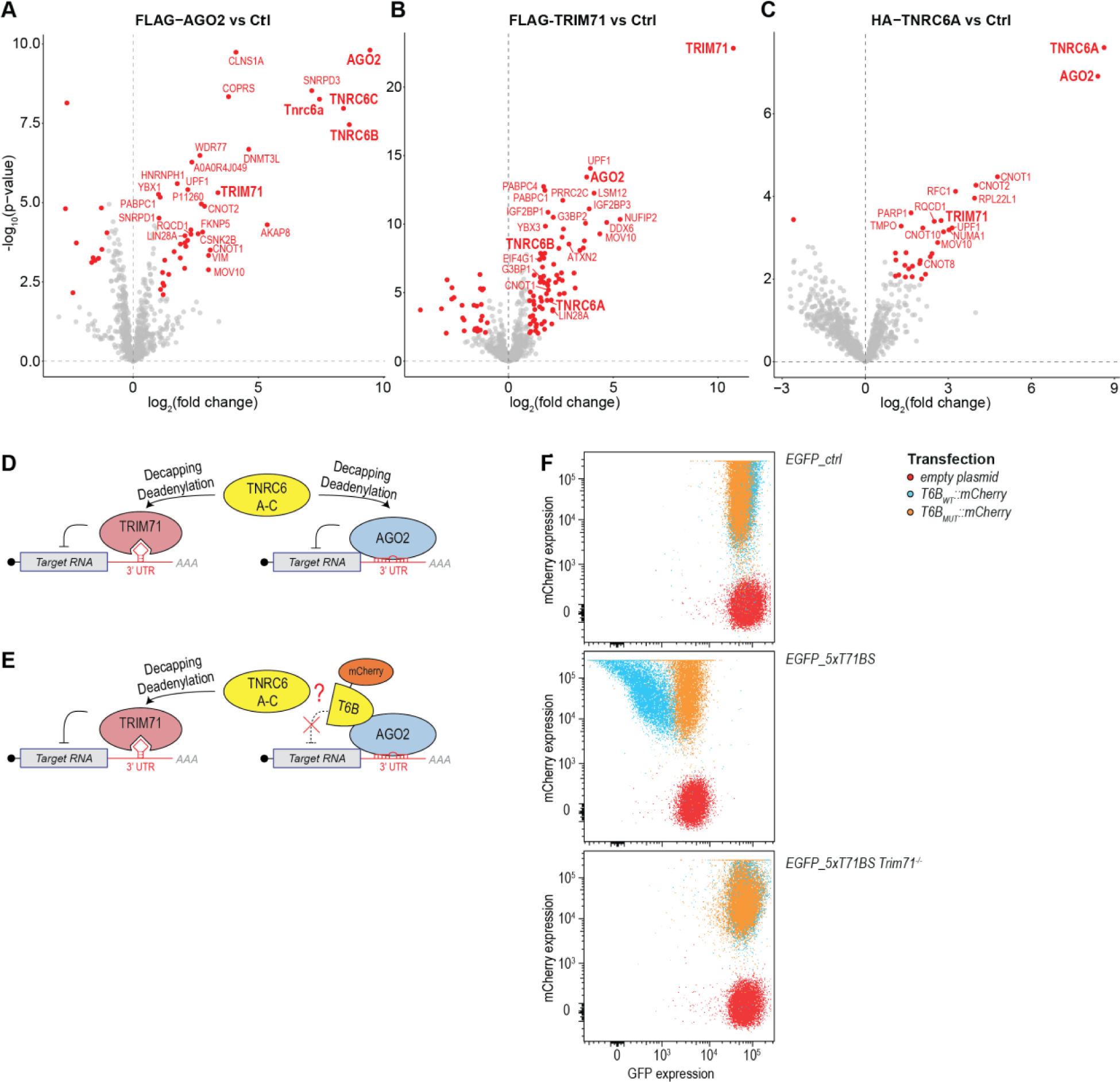
Blocking the interaction between GW182 and AGO2 increases silencing by TRIM71. **(A-C)** Volcano plots showing mass spectrometry results of *FLAG-Ago2* vs untagged control cells (*n* = 3 biological replicates; **[A]**), FLAG-AVI-TRIM71 vs untagged control cells (*n* = 9 biological replicates; **[B]**), and HA-AVI-TNRC6A vs untagged control cells (*n* = 3 biological replicates; **[C]**). Significance was tested using a moderated t-test. Identified proteins with an adjusted *P* value ≤ 0.01 and |log_2_(fold change)| ≥ 1 are marked in red. TNRC6A∼C, AGO2, and TRIM71 are marked bold. Data source **A-C**: **Supplemental Table S2**. **(D)** Schematic drawing showing the working hypothesis of a competition between miRNA, and TRIM71 pathways for TNRC6 as a common silencing factor. **(E)** Schematic drawing illustrating T6B-peptide based competitive inhibition of TNRC6-AGO2 interactions. **(F)** Scatter plots showing FACS results for GFP (x-axis), and mCherry (y-axis) expression in *GFP_Ctrl*, *GFP_5×T71BS*, and *GFP_5×T71BS Trim71^-/-^* reporter cell lines transfected with empty, *T6B_WT_::mCherry*, or *T6B_MUT_::mCherry* plasmids.

We observed reciprocal immunoprecipitation of TRIM71 and AGO2, and both co-immunoprecipitated additional miRNA pathway factors such as DDX6 (**Figure 4A-C**). Moreover, most TRIM71 interactions, including those with AGO2 and TNRC6, were sensitive to treatment with RNase A (**Supplemental Figure S3A, B**). Thus, it seemed possible that physical interactions between TNRC6 and TRIM71 could be mediated by shared targets of miRISC and TRIM71. However, *Ago2* knock-out did not compromise co-immunoprecipitation of TNRC6, or of any other protein interactor with TRIM71 (**Supplemental Figure S3C-E**). Similarly, interaction of TRIM71 with known target RNAs quantified by RNA immunoprecipitation was not altered in *Ago2* knock-out cells (**Supplemental Figure S3F, G**). In conclusion, although TRIM71 – TNRC6 interactions require RNA for stabilization to be captured by immunoprecipitation, they do not rely on mediation by AGO2.

### Blocking the interaction of AGO2 and TNRC6 by peptide-based inhibition results in improved silencing by TRIM71

Given the biochemical and functional interactions between TRIM71 and TNRC6, competition of AGO2 and TRIM71 for TNRC6 appeared to provide a possible explanation for why disruption of miRNA biogenesis enhanced TRIM71 activity (**Figure 4D**). Specific disruption of the TNRC6-AGO2 complex without depletion of either component would provide a stringent test of this model. The interaction of TNRC6 with AGO2 is mediated by GW-amino acid motif repeats in the N-terminus of TNRC6 (ref. (Lazzaretti et al., 2009; Lian et al., 2009; Sheu-Gruttadauria and MacRae, 2018; Takimoto et al., 2009; Zipprich et al., 2009)), permitting displacement of endogenous TNRC6 from AGO2, and inhibition of miRNA activity, through expression of a peptide derived from the HsTNRC6B N-terminus (termed T6B_WT_ in the following)(Hauptmann et al., 2015). We verified that a FH::T6B_WT_::mCherry fusion protein (T6B_WT_), but not a mutant derivative thereof, FH::T6B_MUT_::mCherry (T6B_Mut_), co-immunoprecipitated endogenous AGO2 and AGO1 when stably expressed in mESCs (**Supplemental Figure S4A-D**). However, neither peptide co-immunoprecipitated TRIM71 (**Supplemental Figure S4C, D, Supplemental Table S2**). This suggests that TRIM71 binding to TNRC6 differs from binding by AGO2, allowing us to use *T6B* expression to interfere specifically with the TNRC6–AGO2 complex (**Figure 4E**).

Overexpression of *FH::T6B_WT_::mCherry* but not *FH::T6B_MUT_::mCherry* in the *GFP_5×T71BS* reporter cell line led to a strong and dose-dependent enhancement of repression of the GFP reporter (**Figure 4F**) without changing TRIM71 and AGO2 levels (**Supplemental Figure S4B, E**). This effect was specific, as GFP levels were unaltered in cells lacking TRIM71 or expressing a reporter lacking TRIM71 binding sites **(Figure 4F**).

In summary, displacing TNRC6 from AGO2 enhances the activity of TRIM71, consistent with competition between AGO2 and TRIM71 for limiting amounts of TNRC6 to induce transcript silencing.

### TRIM71 and AGO2 recruit TNRC6A to transcripts that they repress

The physical and functional interactions of TRIM71 and TNRC6 support a model of TNRC6 functioning as a silencing factor for TRIM71-mediated repression. This model predicts recruitment of TNRC6 to TRIM71 targets through TRIM71. To test this model, we performed RNA-immunoprecipitations of endogenously HA-tagged TNRC6A (the choice of isoform for tagging was arbitrary), followed by sequencing (RIP-seq) in wild-type, and *Trim71^-/-^* mESCs. We found that pulldown of 229 genes was significantly decreased in the absence of TRIM71 (**Figure 5A**). Consistent with direct recruitment through TRIM71, TRIM71 CRAC-seq data (Welte et al., 2019) revealed that this set of transcripts was enriched for binding by TRIM71 (**Figure 5B**). Supporting functionality, these transcripts were de-silenced in *Trim71^-/-^* vs WT cells (**Figure 5C, Supplemental Figure S5B**). Finally, an independent set of TRIM71 RIP experiments using RT-qPCR for transcript quantification showed the same trend for the *GFP_5×T71BS* reporter, as well as for the known endogenous TRIM71 target transcripts (*Plxnb2, Mbnl1, Mllt1*; **Supplemental Figure S 5C**). Thus, these data show that TRIM71 recruits TNRC6 to a set of mRNAs that it silences, supporting the notion that TNRC6 is a functional component of a TRIM71-induced silencing complex.

**Figure 5.**
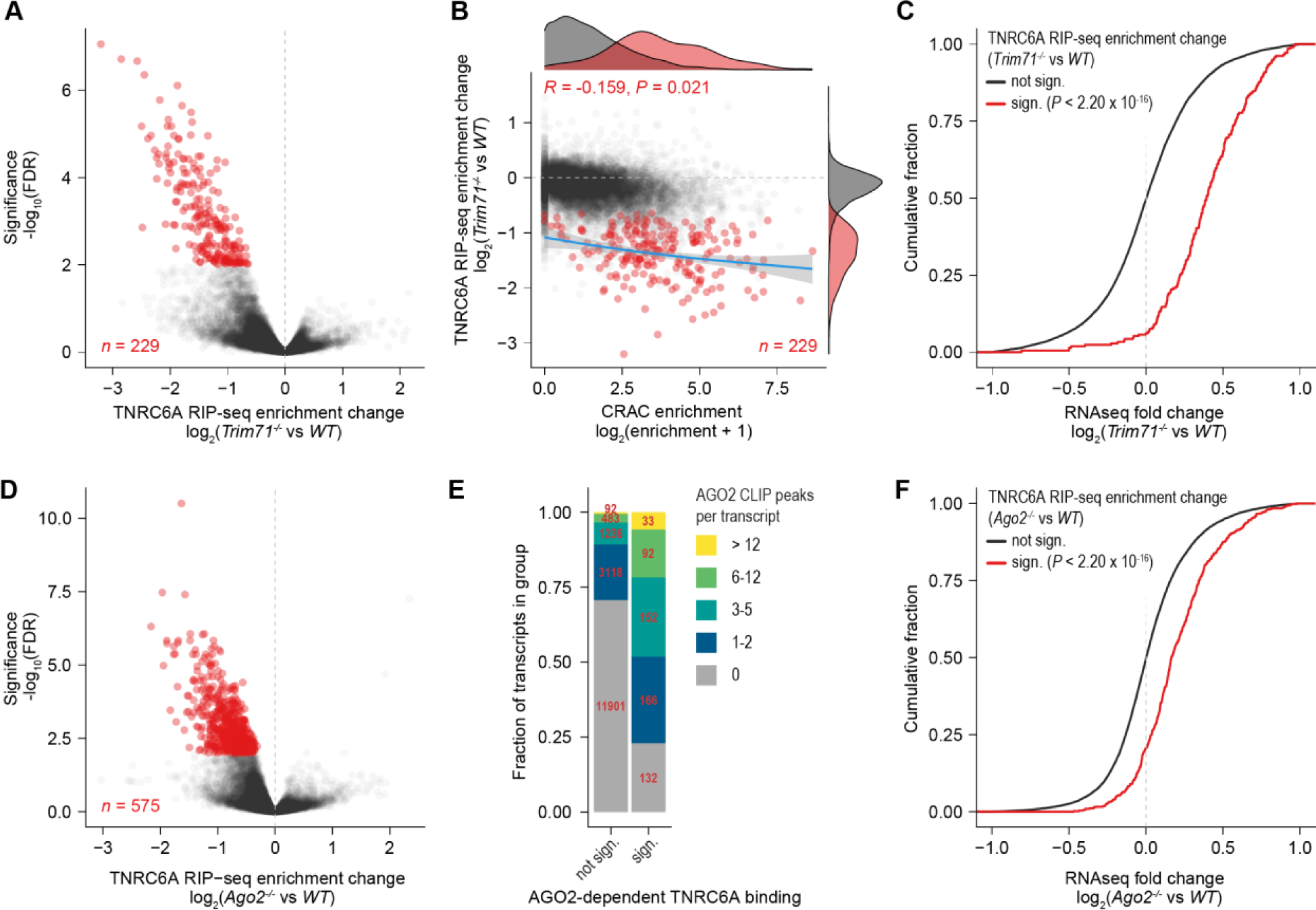
TRIM71 and AGO2 recruit TNRC6A to their respective target RNAs. **(A)** Volcano plot showing changes of TNRC6A transcript binding in *Trim71^-/-^* vs. *WT* cells as quantified by RIP-seq. Significantly depleted transcripts (FDR ≤ 0.01, log_2_[TNRC6A RIP-seq enrichment change] < 0) are indicated in red. **(B)** Scatter plot showing TRIM71 transcript binding assessed by CRAC-seq (x-axis: log_2_[enrichment + 1](Welte et al., 2019)) and TNRC6A RIP-seq enrichment change in *Trim71^-/-^* vs *WT* cells (y-axis). Coloring as in **(A)**. The corresponding density distributions are shown on top for CRAC-seq enrichment and to the right for TNRC6A RIP-seq. *R* indicates Pearson’s correlation coefficient. *P* value is calculated using a paired, two-sided t test for correlation. **(C)** Cumulative density functions showing de-silencing of transcripts with TRIM71-dependent TNRC6A binding (as defined in **[A]**; red line) vs. all other transcripts (black line). Significance was tested with a two-sided Kolmogorov-Smirnov test. **(D)** Volcano plot showing changes of TNRC6A transcript binding in *Ago2^-/-^* vs. *WT* cells as quantified by RIP-seq. Significantly depleted transcripts (FDR ≤ 0.01, log_2_[Tnrc6a RIP-seq enrichment change] < 0) are indicated in red. **(E)** Barplot indicating transcript counts (red numbers) in groups with different numbers of overlapping AGO2 CLIP-seq peaks (Li et al., 2020), as indicated by color labels. Transcripts without (left bar), and with (right bar) significant AGO2-dependent TNRC6A binding (as defined in **[D]**) are separated. **(F)** Cumulative density functions showing de-silencing of transcripts with AGO2-dependent TNRC6A binding (as defined in **[D]**; red line) versus all other transcripts (black line). Significance was tested with a two-sided Kolmogorov-Smirnov test. Data source (**A-F**): **Supplemental Table S3**.

To gain more insight into the interplay of TRIM71- and AGO2-mediated gene regulation, we performed TNRC6A RIP-seq experiments on *Ago2^-/-^* cells (**Supplemental Figure S5A**), along with a repetition of the experiment on wild-type and *Trim71^-/-^* cells. The repeat experiment for *HA-Tnrc6a* and *HA-Tnrc6a Trim71^-/-^* cells replicated the earlier experiment well (**Supplemental Figure S5D**). In *Ago2* knock-out cells, binding of 575 transcripts by TNRC6A was significantly decreased relative to wild-type cells (**Figure 5D**). This set of transcripts was enriched for AGO2 binding events, quantified by CLIP-seq (Li et al., 2020) (**Figure 5E**), and significantly de-silenced in *Ago2^-/-^* relative to wild-type cells (**Figure 5F, Supplemental Figure S5E)**. Most of these transcripts were distinct from those that relied on TRIM71 for TNRC6A binding, *i.e.*, TNRC6A targeting to 43 transcripts depended on both AGO2 and TRIM71, but 532 and 186 target transcripts were unique to AGO2 and TRIM71, respectively (**Supplemental Table S3**). In the following, we will refer to these as ‘common’, ‘AGO2-only’ and ‘TRIM71-only’ targets.

Of note, we observe higher correlation of differential TNRC6 targeting (quantified by RIP-seq) with differential gene expression (quantified by RNA-seq), than with methods quantifying direct transcript binding of TNRC6A-recruiting RBPs (*i.e.*, TRIM71 CRAC-seq or AGO2 CLIP-seq experiments; **Supplemental Figure S5B, E-G**). This either indicates a significant amount of non-functionally relevant binding events, or normalization issues in the latter experimental systems. In summary, AGO2 and TRIM71 recruit TNRC6A to their respective target transcripts, and TNRC6A transcript binding correlates well with silencing activity of the respective pathway.

### Exclusive TRIM71 target transcripts are silenced more efficiently in *Ago2* knock-out cells

The sets of unique and common TRIM71 and AGO2 target transcripts identified above allowed us to ask whether the enhancement of TRIM71 activity upon *Ago2* deletion observed in the genetic reporter screens was true for endogenous TRIM71 targets. Indeed, while ‘TRIM71-only’ and ‘common’ targets were de-silenced in *Trim71^-/-^* relative to wild-type cells (**Figure 6A**), we observed increased silencing of ‘TRIM71-only’ targets in *Ago2^-/-^* relative to wild-type cells (**Figure 6B**). These effects were most pronounced for a subset of transcripts that showed either unchanged, or increased TNRC6A binding in *Ago2^-/-^* relative to WT cells (**Supplemental Figure S6C, D, F, G**). We obtained similar results when comparing gene expression changes between cells expressing *T6B_WT_* and *T6B_MUT_* for these transcript classes (**Figure 6C, Supplemental Figure S6E, H**), consistent with the extensive similarity of gene expression changes introduced by *T6B_WT_* transfection and *Ago2* disruption (**Supplemental Figure S7A**).

**Figure 6.**
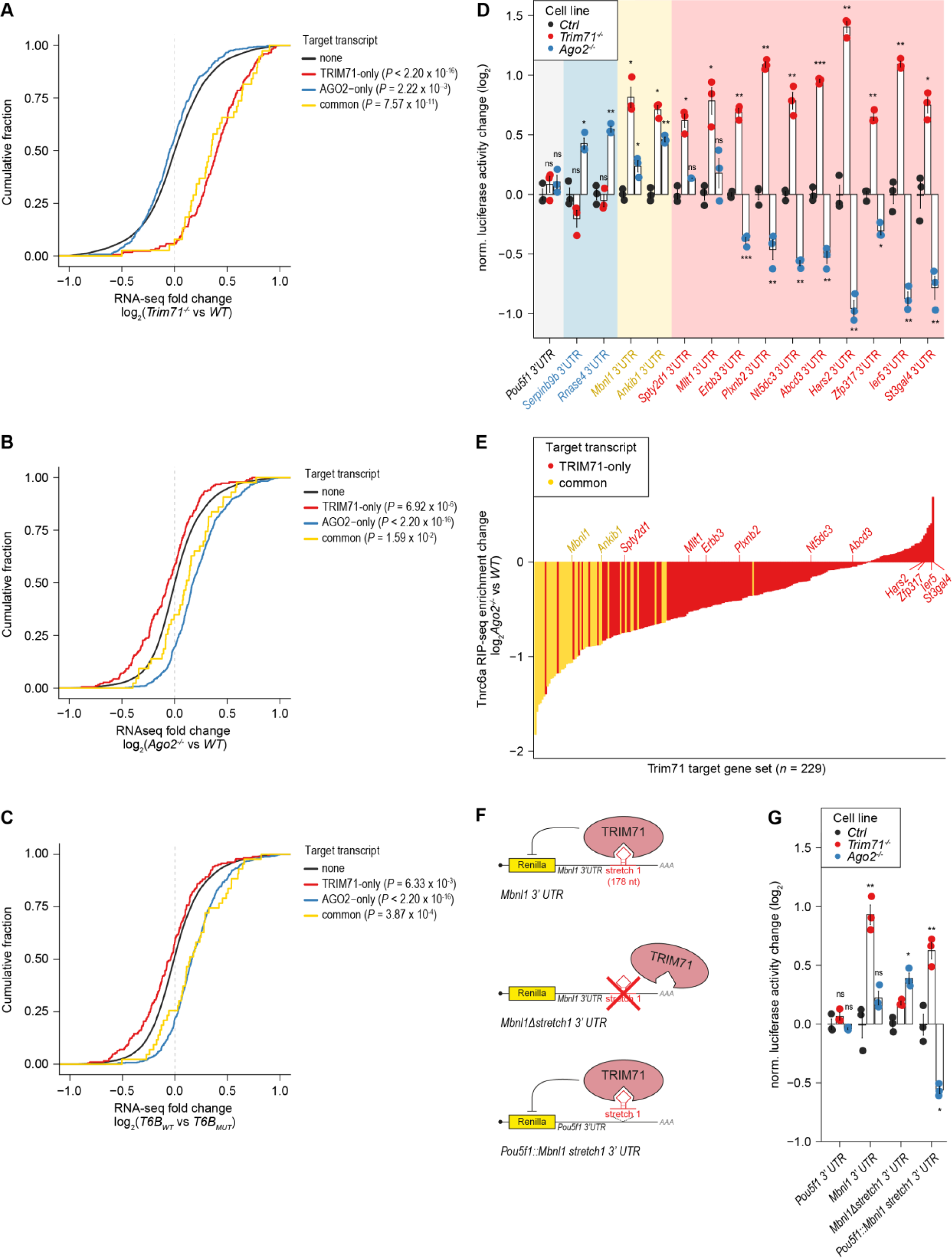
Exclusive TRIM71 target transcripts are silenced more effectively in *Ago2* knock-out cells. **(A-C)** Cumulative density functions showing global expression changes of transcript groups in *Trim71^-/-^* vs *WT* cells **(A)**, *Ago2^-/-^* vs *WT* cells **(B)**, and *T6B_WT_* vs *T6B_MUT_* cells **(C)**. <u>Red color</u> indicates transcripts with significant loss of TNRC6A RIP-seq enrichment in *Trim71^-/-^* vs *WT* cells (FDR ≤ 0.01, log_2_(fold change) < 0), but not in *Ago2^-/-^* vs *WT* cells (*n* = 186). <u>Blue color</u> indicates transcripts with significant loss of RIP-seq enrichment in *Ago2^-/-^* vs *WT* cells (FDR ≤ 0.01, log_2_(fold change) < 0), but not in *Trim71^-/-^*vs *WT* cells (*n* = 532). <u>Yellow color</u> indicates transcripts with significant loss of TNRC6A RIP-seq enrichment in *Trim71^-/-^* vs *WT* cells≤ 0.01, <u>and</u> in *Ago2^-/-^* vs *WT* cells (*n* = 43). <u>Black color</u> indicates transcripts with no significant loss of RIP-seq enrichment in *Trim71^-/-^* vs *WT* cells≤ 0.01, <u>and</u> in *Ago2^-/-^* vs *WT* cells (*n* = 16,509). Significance was tested with a two-sided Kolmogorov-Smirnov test compared to the group ‘none’. **(D, G)** Barplots showing validation experiments with luciferase reporter gene assays with various *Renilla* 3′UTR fusion constructs (selection indicated in **[E** and **Supplemental Figure S7B-G]**) in *WT*, *Trim71* and *Ago2* knock-out mESCs. Background colors indicate grouping of tested transcript, as defined in **(A-C)**. Data were normalized to *WT* and represent mean ± SEM. *n* = 3 biological replicates. Significance was tested using a two-tailed Student’s *t*-test. (*) *P* ≤ 0.05; (**) *P* ≤ 0.01; (***) P < 0.001; (ns) not significant (*P* ≥ 0.05). **(E)** Barplot displaying the ‘TRIM71 target’ gene set as identified in (Figure 5A), ranked by TNRC6A RIP-seq enrichment change in *Ago2^-/-^* vs *WT* cells. Colors indicate grouping of tested transcripts, as defined in **(A-C)**. Transcripts selected for validation experiments with luciferase reporters **(D)** are indicated. **(F)** Schematic drawing showing the different *Mbnl1* 3’UTR reporter constructs used in **(G)**. Background cell line **(A-G)**: *GFP_5×T71BS* mESCs. Data source **(A-C, E)**: **Supplemental Table S3**.

We selected two ’common’, two ‘AGO2-only’ and 10 ‘TRIM71-only’ targets (**Supplemental Figure S7B-G**) to examine their 3’UTR-dependent regulation using luciferase reporters. The ‘TRIM71-only’ set contains two previously validated TRIM71 targets, *Mllt1*, and *Plxnb2* (refs. (Mitschka et al., 2015; Welte et al., 2019)), as well as eight novel reporters. Like the previously validated reporters, all novel TRIM71 target candidates were upregulated in *Trim71^-/-^* relative to WT cells. Conversely, the two ‘AGO2-only’ target candidates were upregulated in *Ago2^-/-^* relative to wild-type cells. This validates AGO2-or TRIM71-dependent recruitment of TNRC6A to a transcript as a powerful criterion for identification of target transcripts for the respective pathway. Indeed, and again consistent with our annotation, expression of the two ‘common’ targets was also increased in *Ago2^-/-^* vs WT cells. Confirming specificity, *Ago2* deletion did not cause upregulation of any of the ‘TRIM71-only’ target reporters. Furthermore, eight of the 10 3’UTRs from ‘TRIM71-only’ targets conferred increased reporter silencing upon *Ago2* knock-out (**Figure 6D, E**), providing additional evidence that miRNA pathway disruption broadly enhances TRIM71-dependent silencing.

To further dissect TRIM71- and miRISC-mediated silencing activities, we analyzed the *Mbnl1* 3’UTR, one of the two ‘common’ 3’UTRs known for both Trim71- and miRNA-mediated silencing (Cerro-Herreros et al., 2018; Welte et al., 2019) (**Figure 6D**). We had previously mapped a functional TRIM71 binding site on this 3’UTR (Welte et al., 2019), allowing us to test whether we could separate the two silencing activities. Indeed, we found that grafting a region (‘stretch1’) that primarily mediates TRIM71-dependent repression (Welte et al., 2019) into the *Pou5f1* 3’UTR conferred de-silencing in *Trim71*^-/-^ but enhanced silencing in *Ago2*^-/-^relative to wild-type cells for the resulting ‘*Pou5f1::Mbnl1 stretch1* 3’UTR’ reporter (**Figure 7F, G**). Conversely, a reporter lacking this TRIM71 binding site (‘*Mbnl1Δstretch1* 3’UTR’) was upregulated in *Ago2*^-/-^ relative to *WT* cells (**Figure 7G**). As observed previously (Welte et al., 2019), regulation of the latter reporter by TRIM71 is blunted but not entirely abrogated, which may lead us to underestimate the extent of its miRNA-mediated regulation.

**Figure 7.**
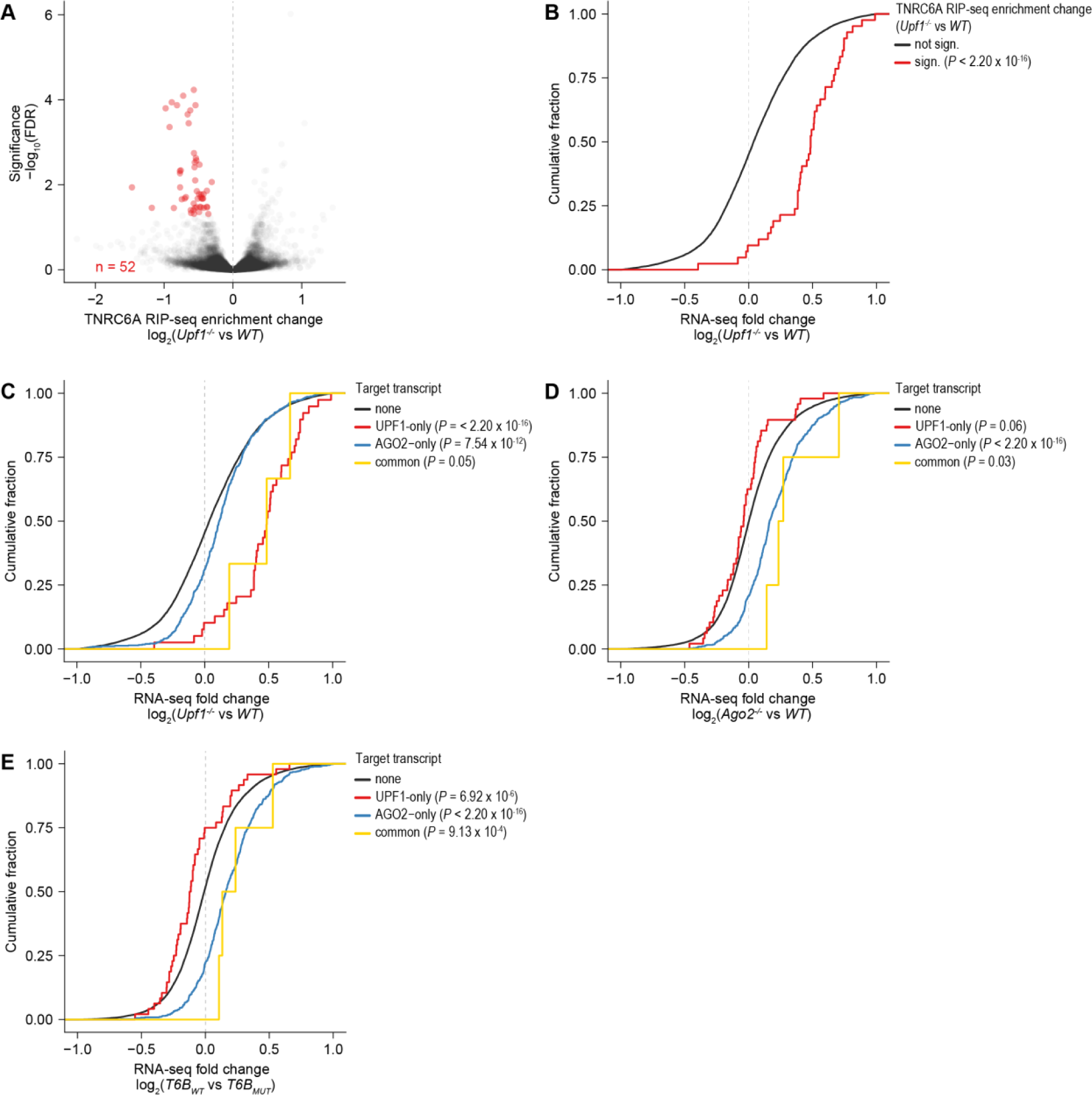
UPF1 recruits TNRC6A to its target RNAs. Exclusive UPF1 target transcripts are silenced more efficiently in *Ago2* knock-out cells. **(A)** Volcano plot showing changes of transcript binding by TNRC6A in *Upf1^-/-^* vs. *WT* cells as quantified by RIP-seq. Significantly depleted transcripts (FDR ≤ 0.05, log_2_[TNRC6A RIP-seq enrichment change] < 0) are indicated in red. **(B)** Cumulative density functions showing de-silencing of transcripts with UPF1-dependent TNRC6A binding (as defined in **[A]**; red line) vs all other transcripts (black line). Significance was tested with a two-sided Kolmogorov-Smirnov test. **(C-E)** Cumulative density functions showing global expression changes of transcript groups in *Upf1^-/-^* vs *WT* cells **(C)**, *Ago2^-/-^* vs *WT* cells **(D)**, and *T6B_WT_* vs *T6B_MUT_* cells **(E)**. <u>Red color</u> indicates transcripts with significant loss of TNRC6A RIP-seq enrichment in *Upf1^-/-^* vs *WT* cells (FDR ≤ 0.05, log_2_(fold change) < 0), but not in *Ago2^-/-^* vs *WT* cells (*n* = 48). <u>Blue color</u> indicates transcripts with significant loss of RIP-seq enrichment in *Ago2^-/-^* vs *WT* cells (FDR ≤ 0.01, log_2_(fold change) < 0), but not in *Trim71^-/-^* vs *WT* cells (*n* = 571). <u>Yellow color</u> indicates transcripts with significant loss of RIP-seq enrichment in *Upf1^-/-^* vs *WT* cells, <u>and</u> in *Ago2^-/-^* vs *WT* cells (*n* = 4). <u>Black color</u> indicates transcripts with no significant loss of RIP-seq enrichment in *Upf1^-/-^* vs *WT* cells, <u>and</u> in *Ago2^-/-^* vs *WT* cells (*n* = 16,648). Significance was tested with a two-sided Kolmogorov-Smirnov test compared to the group ‘none’. Data source: **Supplemental Table S3**.

In conclusion, AGO2 competes with TRIM71 for TNRC6 as a common silencing pathway, and TRIM71’s silencing activity depends on its access to TNRC6.

### Multiple RNA pathways compete with miRISC for access to TNRC6

Finally, we wondered if additional RNA silencing pathways depended on TNRC6. We focused on UPF1, an RNA helicase involved in nonsense-mediated RNA decay as well as many other RNA decay processes (Lavysh and Neu-Yilik, 2020), which reproducibly co-immunoprecipitated with TRIM71, AGO2 and TNRC6 (**Figure 4A-C**) (Elbarbary et al., 2017). To test a potential function of UPF1 in TNRC6 recruitment to specific transcripts, we generated *Upf1^-/-^* cells. Neither the levels of TRIM71 (**Supplemental Figure S7A**) nor silencing of the GFP_5×T71BS reporter were altered (**Supplemental Figure S7B**), showing that UPF1 is not required for regulation of TRIM71 levels or silencing activity.

We performed HA-TNRC6A RIP-Seq in wild-type, *Upf1^-/-^*, and *Ago2^-/-^* cells. The latter served as a positive control and revealed the expected change in TNRC6A recruitment (**Supplemental Figure S7C**). In *Upf1* knock-out cells, binding of 52 transcripts by TNRC6A was significantly decreased relative to wild-type cells (**Figure 7A**). This set of transcripts was significantly de-silenced in *Upf1^-/-^* relative to wild-type cells (**Figure 7B, Supplemental Figure S7D**). Most transcripts were distinct from those that relied on AGO2 for TNRC6A binding, *i.e.*, TNRC6A targeting to only four transcripts depended on both AGO2 and UPF1 (‘common’ targets), but 571 and 48 target transcripts were unique to AGO2 (‘AGO2-only’) and UPF1 (‘UPF1-only’), respectively (**Supplemental Table S3**). Like ‘TRIM71-only’ target transcripts, ‘UPF1-only’ targets displayed increased silencing in *Ago2^-/-^* relative to wild-type and *T6B_WT_* relative to *T6B_MUT_* cells (**Figure 7C-E, Supplemental Figure S7E-J**). We conclude that UPF1 recruits TNRC6 to a subset of its targets to achieve silencing, and that this silencing activity is enhanced in the absence of AGO2. In summary, our data reveal that AGO2 can influence transcripts directly, through miRNA-guided silencing, and indirectly, by competing TNRC6 away from multiple RNA pathways (*i.e.*, TRIM71 and UPF1), thus constraining their silencing activity.

## Discussion

TNRC6/GW182 has been known as a core component of miRISC (Braun et al., 2011; Chen et al., 2009; Jakymiw et al., 2005; Liu et al., 2005). Our results extend this canonical function by revealing TNRC6 silencing activities in additional pathways, depending on distinct RBPs. A shared dependence on TNRC6 provides a mechanism for cross-regulation among these pathways and allows AGO2 to constrain the activities of additional RNA silencing pathways by sequestering TNRC6. As TRIM71 in turn can repress production of certain miRNAs(Torres-Fernández et al., 2021b; Treiber et al., 2017) and of AGO2 (ref. (Liu et al., 2021)), a cross-regulatory network of interactions among RNA silencing pathways is beginning to emerge.

### TNRC6 mediates mRNA silencing by TRIM71

Several lines of evidence support the conclusion that TNRC6 has a critical function in mediating TRIM71-dependent RNA silencing: First, over-expression of *TNRC6* paralogs enhances TRIM71 activity. Second, TNRC6 co-immunoprecipitates with TRIM71. Although this interaction is RNA-dependent, it does not simply reflect a coincidental co-occurrence on the same transcript, as third, we find that TRIM71 mediates recruitment of TNRC6 to specific mRNAs. Accordingly, co-immunoprecipitation of TNRC6 with TRIM71 does not depend on the presence of the known TNRC6 binding partner AGO2. Fourth, transcripts that rely on TRIM71 or AGO2 for TNRC6 recruitment are functional targets of each pathway, and largely fall into distinct sets.

Strikingly, loss of TNRC6 binding upon TRIM71 or AGO2 inactivation appears to be a better predictor of each pathway’s silencing activity than direct transcript binding of TRIM71 or AGO2 (**Supplemental Figure S5B, E-G**). Thus, the presence of TNRC6 can distinguish productive from non-productive binding events of either RBP. The fact that co-immunoprecipitation of TNRC6 with TRIM71 is sensitive to RNase treatment further suggests that RNA-binding of TRIM71 may support recruitment of, or stabilize the interaction with, TNRC6.

We note that weak predictive power of RNA binding for function is not unique to TRIM71 or AGO2, but a general challenge in experimental systems seeking to quantify direct RBP-RNA interactions (Chen et al., 2019; Vieira-Vieira and Selbach, 2021). Hence, differential recruitment of silencing effectors – TNRC6 as shown here, but potentially also others for other RBPs – appears to be a robust measure to identify functional targets in high throughput and, potentially, diverse contexts.

### TNRC6 functions outside the miRNA pathway

Although mESCs with deletions of miRNA pathway components *(Ago2, Dicer, Xpo5*) are viable, we were unable to obtain *Tnrc6a-c* triple deletion cells, consistent with functions of TNRC6 beyond miRISC. As cells lacking either *Trim71* or combinations of *Trim71* and miRNA pathway genes are also viable, these ‘non-canonical’ TNRC6 functions appear not to be restricted to TRIM71. Our demonstration that UPF1 also recruits TNRC6 to specific transcripts for silencing supports this notion. We emphasize that UPF1 causes dysregulation of many transcripts (*n* = 8,074; FDR-cutoff: ≤ 0.05), among which only a small subset exhibit altered TNRC6A recruitment. While we have not investigated which features route specific UPF1 targets to this particular decay pathway, the finding is consistent with the fact that UPF1 employs diverse mechanisms to achieve transcript silencing, including but not limited to its well-established function in NMD(Cho et al., 2015; Choe et al., 2014; Kaygun and Marzluff, 2005; Kim et al., 2005; Kim and Maquat, 2019; Lavysh and Neu-Yilik, 2020; Mino et al., 2015; Park et al., 2016). Irrespectively, our findings demonstrate that TNRC6’s function as an RNA silencing factor extends well beyond its canonical role in the miRNA pathway.

### The interplay of miRNA, TRIM71 and UPF1 pathways on mRNA silencing

Through various manipulations, we have shown that TRIM71 and UPF1 activity is enhanced when TNRC6 is released from binding by AGO2. The opposite effect, enhanced miRNA activity upon loss of TRIM71 or UPF1, appears weak at best. This may reflect a higher affinity of TNRC6 for AGO2 so that AGO2 can sequester TNRC6 away from other RBPs, while these cannot, or much less efficiently, sequester TNRC6 away from AGO2. Indeed, TNRC6 forms stable complexes with AGO2 in the absence of RNA (Elkayam et al., 2017; Hicks et al., 2017; Sheu-Gruttadauria and MacRae, 2018), distinguishing them from the complexes formed with TRIM71 and UPF1. The fact that the T6B peptide blocks binding of TNRC6 to AGO but enhances TRIM71 and UPF1 activity further indicates that TNRC6 can utilize distinct domains for binding to different RBPs, which future structure-function studies may identify.

Although the main effect of miRNA pathway inactivation is an enhancement of TRIM71 and UPF1 silencing activities by increasing TNRC6 availability, more complex responses result for transcripts under simultaneous regulation by AGO2 and either RBP. Incidentally, this validates the benefit of performing a screen with a heterologous 3’UTR to isolate the regulatory phenomenon under investigation from complex cellular regulatory circuits and their confounding effects. Thus, for *Mbnl1*, we showed that its 3’UTR mediates regulation by both AGO2 and TRIM71. This leads to the paradox situation that miRNA pathway inactivation does not promote transcript accumulation, because enhanced TRIM71-dependent silencing compensates for a loss of miRNA-mediated silencing.

Additional scenarios are conceivable: Appropriately spaced miRNA-binding sites can cooperatively increase transcript silencing through a mechanism that occurs downstream of target binding by the miRNA (Broderick et al., 2011; de la Mata et al., 2015), possibly at the level of TNRC6 binding (Elkayam et al., 2017). It would thus seem possible that cross-pathway synergy could also occur among appropriately spaced miRNA-, TRIM71-, and UPF1-binding sites. Although not investigated here, support for such a mechanism comes from previous work on the *Cdkn1a* mRNA, which suggested cooperation of TRIM71 with miRNAs and UPF1, to achieve silencing of this particular transcript in mouse and human cells, respectively (Chang et al., 2012; Torres-Fernández et al., 2019). Nonetheless, such cooperative pathway activity - at least in mESCs - appears to be the exception, as we observe little overlap among the target sets of TRIM71, AGO2 and UPF1, respectively.

## Acknowledgements

We are grateful to Kathrin Braun and Christiane Wirbelauer for help with cell culture work. We thank Sebastien Smallwood, Eliza Pandini Figueiredo Moreno, Stéphane Thiry, Alicia Lindeman and Walter Carbone for library preparations and high-throughput sequencing and Hubertus Kohler for FACS-sorting. We thank Ulrich Elling for input on CRISPR/sgRNA-screening experiments, and Gunter Meister for T6B plasmids. We thank Rajani Gudipatti and Florian Aeschiman for fruitful discussions and ideas for experiments, and Marc Bühler, Dimosthenis Gaidatzis, and Witold Filipowicz for critically reading manuscript drafts. This project has received funding from the Swiss National Science Foundation through the NCCR RNA & Disease (Grant No. 182880 and 205601, to H.G.), and the Novartis Research Foundation through Friedrich Miescher Institute for Biomedical Research core funding (to H.G.). The work of T.W. was funded by the Deutsche Forschungsgemeinschaft (DFG, German Research Foundation), Grant No. 466417053. M.J.L. was supported by a predoctoral fellowship from the Boehringer Ingelheim Fonds.

## Author Contributions

T.W. conceived the project, performed molecular cloning, genome edited cell lines, designed and performed experiments, computationally analyzed sgRNA screens, RNA-seq, RIP-seq, mass-spectrometry results, and wrote the manuscript. A.N., A.G., C.A., and M.J.L. created plasmid constructs and cell lines, and performed experiments. M.B.S. computationally analyzed sgRNA screens, RNA-seq, and RIP-seq experiments. D.Hess performed and computationally analyzed mass-spectrometry experiments. J.S., and C.S. developed mass-spectrometry analysis software. I.S. performed FACS-sorting for sgRNA-screens. F.N. performed initial bioinformatic analysis of sgRNA screens. D.Hoepfner., D.E., and J.R-H. provided technical support and reagents. H.G. conceived the project, designed and analyzed experiments, and wrote the manuscript.

## STAR Methods

### Materials Availability

Helge Großhans, Friedrich Miescher Institute for Biomedical Research, will respond to requests and provide reagents and information.

Published research reagents from the FMI are shared with the academic community under a Material Transfer Agreement (MTA) having terms and conditions corresponding to those of the UBMTA (Uniform Biological Material Transfer Agreement).

### Data and Code Availability

All the data is included in the manuscript or in the supplementary material. Next-generation sequencing data have been deposited at the Gene Expression Omnibus with accession numbers GSE205968 (https://www.ncbi.nlm.nih.gov/geo/query/acc.cgi?acc=GSE205968) and GSE214755 (https://www.ncbi.nlm.nih.gov/geo/query/acc.cgi?acc=GSE214755). The mass spectrometry proteomics data have been deposited to the ProteomeXchange Consortium via the PRIDE partner repository (Perez-Riverol et al., 2022) with the dataset identifier PXD034537 (http://www.ebi.ac.uk/pride). The einprot R package (v0.4.4) used for the analysis of mass spectrometry data is deposited online (https://github.com/fmicompbio/einprot).

## Experimental Model and Subject Details

### Mouse embryonic stem cells (mESCs)

mESCs (TC-1 mouse ES cells (background 129S6/SvEvTac, a gift from D. Schübeler (Lienert et al., 2011) and 129xC57BL/6 *Rosa26^Cre-ERT2/BirA-V5^* [cMB063]) (Ostapcuk et al., 2018) were cultured on gelatin-coated dishes in mESC medium (DMEM [Gibco 21969-035] supplemented with 15% fetal bovine serum [Gibco], 1 × non-essential amino acids [Gibco], 1 mM sodium pyruvate [Gibco], 2 mM L-glutamine [Gibco], 0.1 mM ß-mercaptoethanol [Sigma], 50 μg/ml penicillin, 80 μg/ml streptomycin and home-made LIF) at 37°C in 5% CO_2_.

## Method Details

### Generation of cell lines

Generated cell lines can be found in **Supplemental Table S5**. Experimental details for the generation of each cell line are given in the **Supplemental Methods** section.

### Molecular cloning

Generated plasmids with insert sources and primer sequences can be found in **Tables S6 and S7**. Experimental details for the generation of plasmid constructs are given in the **Supplemental Methods** section.

### Genome-wide sgRNA screen

Genome-wide sgRNA-screening was performed as described previously (Hoepfner et al., 2019). Briefly, each cell line (mESC *Cas9::BSD*, yGlob [*GFP_Ctrl*; *GFP_5×T71BS*]) was expanded to confluency in 8 x 15 cm plates in mESC-medium containing 2.5 µg/ml Blasticidin. Cells were transduced with two separate lentiviral sgRNA libraries (library 1: 52,000 unique sgRNAs (∼5 per gene); library 2: 39,000 unique sgRNAs (∼5 per gene); (Zeng et al., 2018). Cells were transduced at a multiplicity of infection (MOI) of 0.31 using 10 µg/ml polybrene at 700× coverage/sgRNA. In total, 126.25×10^6^ cells were transduced with library 1 on eight T300 plates [TPP, #90301] and 94.7x10^6^ cells were transduced with library 2 on six T300 plates. Transduced cells were selected with Puromycin at 1 µg/ml for three days. On day 8, cells were trypsinized, PFA-fixed (4% PFA in PBS, 15 min at room temperature), washed in PBS, and resuspended in FACS-buffer (2% FBS, 2 mM EDTA in PBS), and FACS-sorted to obtain ‘high 25%’ and ‘low 25%’ signal fractions per cell line. At day 4 and 8, unsorted fractions (200×10^6^ cells per library 1 and 150×10^6^ cells per library 2) were collected. Genomic DNA was extracted, followed by library preparation and sequencing as described previously (Hoepfner et al., 2019).

### Validation sgRNA-Screen

Using the top 985 enriched and depleted genes (based on a z-score ranking sgRNAs according to their effect on GFP expression specific to the TRIM71 pathway; see bioinformatic methods in the **Supplemental Methods** section), respectively, as well as 400 middle-scoring genes from the genome-wide sgRNA screen, we constructed a 12,000 sgRNA library (∼5 sgRNAs per gene) using a previously published sgRNA library (www.vbc-score.org; (Michlits et al., 2020)). sgRNA sequences can be found in **Supplemental Table S1**. The setup for the validation screen corresponded to the whole-genome screen with appropriately down-scaled cell numbers. Due to low viral titers of the library, cells were transfected at a MOI of 0.1, with a 500-fold coverage.

### Experiments with transiently transfected mESCs

5.0×10^5^ cells were seeded in 6-well plates and transfected with 2 µg of plasmid using Lipofectamine 3000 [Invitrogen; L3000] reagent at 1 μl/μg ratio in OptiMem medium [Gibco; 31985062]. After 24h of transfection, cells transfected with Puromycin-containing constructs were selected with puromycin at 1 μg/ml for two days. For FH-TNRC6A-C constructs, puromycin concentrations were reduced by 50% at day two.

### Luciferase assays

Luciferase assays were performed as described previously (Welte et al., 2019). Briefly, 25,000 cells were transfected with 100 ng of plasmid DNA in 24-well plates using Lipofectamine 3000 (Invitrogen, L3000001) according to manufacturer’s instructions. psiCHECK-2™ (Promega, C8021) luciferase plasmids and primers used can be found in **Supplemental Table S6B** and **Supplemental Table S7C**. Transfections were carried out in triplicates. The following day, cells were washed once with PBS, lysed, and Renilla and Firefly luciferase activities were measured on a luminometer using the Promega dual luciferase kit (Promega, E1910) according to manufacturer’s instructions.

### RNA extraction, RT-qPCR and RNA-seq experiments

RNA extraction, RT-qPCR, and RNA-seq was done as previously described **(Welte et al., 2019)**. Briefly, total RNA was extracted using the Norgen single-cell RNA purification kit (Norgen, 51800) with on-column DNase digestion (Norgen, 25710).

For qPCRs, 400 ng of total RNA was transcribed to cDNA using the ImProm-II™ reverse transcription kit (Promega, A3500) with random primers. qPCRs were performed using the SYBR™ Green PCR master mix (Applied Biosystems, 4309155) with standard cycling conditions. Data were normalized to *Gapdh* expression levels. qPCR primer sequences can be found in **Supplemental Table S8**.

Sequencing library preparation for RNA-seq was done using the TruSeq™ 2 (Illumina, RS-122-2001) kit, followed by HiSeq2500 (RNA-seq experiments from **Figs. 5-7, Extended Data Figs. S5-S7**), or SMART-Seq™ 2 (Illumina, 20040532), followed by NovaSeq (RIP-seq experiments from **Figs. 5-7, Extended Data Figs. S5-S7**).

### Western Blotting

Western Blotting was performed as described previously (Welte et al., 2019). Briefly, cells were seeded at equal density in 6-well plates. The following day, cells were lysed with lysis buffer (25 mM Tris-HCl at pH 7.5, 100 mM KCl, 0.5% [v/v] NP-40, 1 mM EDTA, EDTA-free cOmplete mini protease inhibitor [Roche, 1186170001]). 40 µg of protein was loaded onto NuPAGE-Novex Bis-Tris 4%–12% gradient gels (Invitrogen), transferred semidry to PVDF membranes, blocked for 1 h with 5% milk in TBS plus 0.1% Tween 20 (TBST), and stained with primary antibodies. Antibodies, dilutions and incubation conditions are summarized in **Supplemental Table S9**.

### Immunoprecipitations with endogenously tagged proteins

Immunoprecipitations were performed as described previously (Welte et al., 2019) with the following changes: Cells were grown to confluency in 10-cm dishes. Cells were washed 1x with PBS and lysed with lysis buffer (25 mM Tris-HCl at pH 7.5, 100 mM KCl, 0.5% [v/v] NP-40, 1 mM EDTA, EDTA-free cOmplete mini protease inhibitor [Roche, 1186170001], PhosSTOP [Roche, PHOSS-RO]. If indicated, RNaseA (250 µg/ml; Quiagen, 19101), or RNase inhibitors [SUPERase•In™ RNase Inhibitor, Invitrogen, AM2694] were added. Lysates were cleared by centrifugation. Equal amounts of protein were incubated with 20 µL of FLAG M2 magnetic beads (Sigma, M8823), or Pierce™ HA magnetic beads (Thermo Scientific, 88836) for 2 h at 4°C. Beads were washed three times with 200 µL of washing buffer (50 mM Tris-HCl at pH 7.5, 150 mM KCl, 5 mM MgCl_2_, 0.05% NP-40, EDTA-free cOmplete mini protease inhibitor [Roche, 11836153001], PhosSTOP [Roche, PHOSS-RO]). If indicated, RNaseA (250 µg/ml; Quiagen, 19101), or RNase inhibitors [SUPERase•In™ RNase Inhibitor, Invitrogen, AM2694] were added. For RIP-seq or qPCR, pulldown RNA was isolated as described above. For mass spectrometry, beads were washed once with 70 µl washing buffer without NP-40.

### Mass Spectrometry

On-bead digestion of proteins was performed using 0.2 μg LysC [Wako Chemicals] for 6 h at 37 °C in 2.5 M guanidine-HCl, containing 20 mM 3-[4-(2-hydroxyethyl)-1-piperazinyl]-1-propanesulfonic acid (EPPS) pH 8.5, 10 mM chloroacetamide (CAA) and 5 mM Tris(2-carboxyethyl)phosphine hydrochloride (TCEP) in a total volume of 6 μl. Samples were diluted with 18 μl 50mM HEPES pH 8.5 containing 0.2 μg modified porcine trypsin (Promega) and cleaved overnight at 37 °C. Another 0.2 μg trypsin was added and the cleavage was continued for 6 h.

The generated peptides were acidified with 0.8% TFA (final concentration) and analyzed by LC–MS/MS on an EASY-nLC 1000 [Thermo Scientific] with a two-column set-up. The peptides were applied onto a peptide μPACᵀᴹ trapping column in 0.1% formic acid, 2% acetonitrile in H2O at a constant flow rate of 5 μl/min. Using a flow rate of 500 nl/min, peptides were separated at RT with a linear gradient of 3%–6% buffer B in buffer A in 4 min followed by a linear increase from 6 to 22% in 55 min, 22%–40% in 4 min, 40%–80% in 1 min, and the column was finally washed for 13 min at 80% buffer B in buffer A (buffer A: 0.1% formic acid; buffer B: 0.1% formic acid in acetonitrile) on a 50 cm μPACᵀᴹ column [PharmaFluidics] mounted on an EASY-Spray™ source [Thermo Scientific] connected to an Orbitrap Fusion LUMOS [Thermo Scientific]. The data were acquired using 120,000 resolution for the peptide measurements in the Orbitrap and a top T (3 s) method with HCD fragmentation for each precursor and fragment measurement in the ion trap according the recommendation of the manufacturer (Thermo Scientific).

### Staining of CAS9 for Flow cytometry

Staining of FLAG-CAS9 was done as follows: 1.5×10^6^ cells were PFA-fixed in 4% PFA (Pierce™ [Thermo Scientific, 28908]) for 10 min at room temperature. Cells were washed once in PBS, followed by incubation with blocking solution (1% BSA, 3% Donkey Serum [Sigma: D9663], 0.1% (v/v) Tween in PBS) for 1-2 hours at room temperature. Cells were spun down and resuspended in blocking solution with 1:500 anti-FLAG M2 monoclonal antibody (Sigma [F1804]) and incubated over night at 4°C. Cells were washed twice with flow cytometry solubilisation buffer (R&D [FC005]) + 0.1% BSA, followed by incubation for 45 min at room temperature with Goat Anti-Mouse Alexafluor 647 secondary antibody (Thermo Scientific, A-21235) in flow cytometry solubilisation buffer + 0.1% BSA. Cells were washed twice with flow cytometry solubilisation buffer, followed by flow cytometry.

### FLAER assay

A FLAER (Fluorescently, Alexa488-labeled, inactive variant of aerolysin) assay to quantify CAS9 editing efficacy was performed as described previously (Hoepfner et al., 2019). At day 7, cells were stained with FLAER reagent (Alexa 488 proaerolysin variant [Cedarlanelabs, FL2S]), and analyzed by flow cytometry.

### Flow cytometry

Cells were trypsinized, spun down, and resuspended in PBS. Flow cytometry analysis was done using a BD LSRII SORP analyzer. Unless stated otherwise a stop-gate count of 10,000 events was used. For FACS sorting, resuspended cells were filtered through CellTrics 30 µm filter tubes [Wolflabs; 040042326], followed by sorting using MA900 [Sony], FACSAria [BD], or Influx [BD] cell sorters. Analysis of FACS data was performed using FlowJo 10.8.1 software.

## Quantification and Statistical Analyses

### CRISPR-Cas9 screen data analysis, RNA-seq data analysis, mass spectrometry analysis

A detailed description is given in the **Supplementary Methods** section.

### RT-qPCR analysis, luciferase assay analysis

RT-qPCR and luciferase assays are three biological replicates. The value of each replicate is indicated with a dot. Error bars indicate the mean standard error SEM. *P*-values are calculated using a two-tailed Student’s T test.

## Supplemental Material

Supplemental Methods

Supplemental Figure S1 – 10

Legends for Supplemental Table S1 – 3 (Excel files not included in this document)

Supplemental Table S4 – 9

Supplemental References

## Supplemental Methods

### Generation of cell lines

mESC Cas9::BSD:

3×10^6^ TC-1 mESCs (background 129S6/SvEvTac, gift from D. Schübeler with a recombinase-mediated cassette exchange (RMCE) site located in the gamma-globin gene locus(Lienert et al., 2011) were transfected with 3 µg of pC2P *Cas9::BSD* (pTW71, see **Supplemental Table S6A**), followed by Blasticidin selection. Transfections were carried out using Lipofectamine 2000 [Invitrogen; L2000] reagent at 3.3 μl/μg ratio in Opti-MEM medium [Gibco; 31985062]. Single clones were isolated and tested for Cas9 expression by Western Blot and Cas9 staining. The optimal titer for viral transfection was determined using a reference virus expressing RFP. Cas9 editing capability verified via a FLAER-Assay according to manufacturers’ instructions(Brodsky, 2009) Briefly, 50,000 cells were plated, infected with either gRNA targeting the PIG-A gene or control virus in ESC media containing 10 μg/ml polybrene on day one, and were selected with Puromycin at 1 µg/ml for three days from day two. On day seven, 200,000 suspended cells were stained with 1:100 FLAER reagent in 3 % BSA, fixed in 4% fomaldehyde in PBS, and the ratio of FLAER-negative cells was determined by FACS direcly correlating with Cas9 mediated editing (**Supplemental Figure S1A-C**).

mESC Cas9::BSD, yGlob (GFP_Ctrl & GFP_5×T71BS):

These cell lines were generated using RMCE as described previously(Lienert et al., 2011; Yang et al., 2006), using pL1_1L_pCAG_GFP::unc54 3UTR (‘GFP_Ctrl’; pTW97) & pL1_1L_pCAG_GFP::unc54::mab10_condensed_ 3UTR (‘GFP_5×T71BS’; pTW98).

*mESC Cas9::BSD, yGlob (GFP_Ctrl & GFP_5×T71BS) Trim71^-/-^, Ago^-/-^, Dicer^-/-^, Xpo5^-/-^:* 5×10^5^ *mESC Cas9::BSD, yGlob (GFP_Ctrl & GFP_5×T71BS)* were seeded per well in 6-well plates. Cells were transfected with 2 µg of respective pC2P sgRNA plasmids (sgRNAs: see **Supplemental Table S4**). After 24 h of transfection, cells were selected with Puromycin at 1 μg/ml for 50 hours. Knockout was confirmed by Western Blotting.

*mESC Cas9::BSD, yGlob (GFP_5×T71BS), HA-AVI-Tnrc6a:*

5×10^5^ cells were transfected with 1 µg each of pC2P Tnrc6a_sgRNA-Nterm and ssODN_Tnrc6a_HAAVI (see **Supplemental Table S5**). Transfected cells were selected by Puromycin at 1 μg/ml 24 h after transfection, followed by clonal selection and genotyping (Primers: see **Supplemental Table S7A**).

*mESC Cas9::BSD, yGlob (GFP_5×T71BS), HA-AVI-Tnrc6a Trim71^-/-^, Ago^-/-^:*

5×10^5^ *mESC Cas9::BSD, yGlob (GFP_5xT71BS), HA-AVI-Tnrc6a* were seeded per well in 6-well plates. Cells were transfected with 2 µg of respective pC2P sgRNA plasmids (sgRNAs: see **Supplemental Table S4**). After 24 h of transfection, cells were selected with Puromycin at 1 μg/ml for 50 hours. Knockout was confirmed by Western Blotting.

*mESCCas9::BSD, yGlob (GFP_5xT71BS) FLAG-HA-T6B_WT_::mCherry,mESC Cas9::BSD, yGlob (GFP_5xT71BS) FLAG-HA-T6B_MUT_::mCherry:*

5x10^5^ cells were transfected with 2 µg each of pCIP FH-T6B_WT_::mCherry (pTW114), pCIP FH-T6B_MUT_::mCherry (pTW115). Transfected cells were selected by Puromycin at 1 μg/ml 24 h after transfection for 7 days, followed by one round of FACS-sorting (for mCherry fusion proteins) to exclude mCherry-negative cells. Expression of constructs was evaluated by Western Blot.

### Molecular cloning

Primer sequences for the generation of the respective plasmid number can be found in **Supplemental Table S7B, S7C.**

*Generation of pC2P Cas9::BSD (pTW71):*

pC2P *Cas9::BSD* (pTW71) was created from pTW42(Welte et al., 2019). Deletion of the U6 promoter plus sgRNA scaffold was achieved by digestion with *PcI*I, *Kpn*I, blunting and ligation. Exchange of Puromycin to Blasticidin was achieved by digestion with *Spe*I, *Not*I. Blasticidin was amplified from pEF1-B (Matyas Flemr). BGHPA was amplified from pC2P (pTW71). Fragments were assembled by Gibson assembly.

*Generation of pL1_1L_CAG_GFP::unc54 3UTR (‘GFP_Ctrl’, pTW97) and pL1_1L_CAG_GFP::unc54::mab10_condensed_ 3UTR (‘GFP_5xT71BS’, pTW98):* pTW47 (*GFP_Ctrl*) and pTW48 (*GFP_5xT71BS*)(Welte et al., 2019) were digested with *Spe*I/*Pac*I to cut out the expression cassette (Promoter, GFP, 3UTR, poly-A sequence). pL1_1L was amplified from pL1_LPP-1_1L(Lienert et al., 2011). Fragments were assembled by Gibson assembly.

*pC2P Ago2, Dicer1, Xpo5, Tnrc6a sgRNA plasmids:*

Plasmids were generated by cloning sgRNAs into pC2P as described previously(Welte et al., 2019). A list of sgRNAs can be found in **Supplemental Table S4**.

*Generation of pCIP Ago2 (pTW110), pCIP FH-HsTNRC6A (pTW111), pCIP FH-HsTNRC6C (pTW113):*

MmuAgo2, FH-HsTNRC6A, and FH-HsTNR6C were amplified from Addgene plasmids #73538, #19883, #19885, respectively. Fragments were Gibson-assembled into the *BamHI* site of pCIP (Addgene #79009).

*Generation of pCIP HA-HsTNRC6B (pTW112)*:

HsTNRC6B was amplified from Addgene plasmid #10979. The FH-Tag was amplified from pCIP FH-HsTNRC6A (pTW111). Fragments were Gibson-assembled into the *BamHI* site of pCIP (Addgene #79009).

*Generation of pCIP FH_T6B_WT__mCherry and pCIP FH_T6B_MUT__mCherry (pTW114, pTW115):*

Plasmids were generated by cloning FH_T6B_eYFP and FH_T6B_mut_eYFP(Hauptmann et al., 2015) into the BamHI restriction site of pCIP by Gibson assembly. EYFP from FH_T6B_eYFP and FH_T6B_mut_eYFP was replaced with mCherry by digestion with *Pac*I and *Not*I. mCherry was amplified from pTW51(Welte et al., 2019). Fragments were Gibson-assembled.

*Luciferase reporter gene constructs:*

psiCHECK-2 Abcd3_3’UTR (pTW132), Nt5cd3_3’UTR (pTW133), Spty2d1_3’UTR (pTW134), Ankib1_3’UTR (pTW135), Hars2_3’UTR (pTW136), St3gal4_3’UTR (pTW137), Ier5_3’UTR (pTW138), Erbb3_3’UTR (pTW139): 3’ UTRs were amplified from mESC genomic DNA and Gibson-assembled into the NotI-site of psiCHECK-2.

### Bioinformatic analyses

Bioinformatic analyses and data visualization were performed in R v4.1.1. and v.4.1.2 (R Core Team, 2021).

#### sgRNA screens

SgRNA counts were generated by aligning raw sequences to the reference libraries using Bowtie(Langmead et al., 2009), allowing for no mismatches. Normalization and differential fold change estimates between samples for each sgRNA were generated using edgeR(Robinson et al., 2010), version 3.28.0, using the functions *calcNormFactors* with raw sgRNA counts and default parameters as inputs and *cpm* with parameters “log = TRUE, prior.count = 4.0”. Log_2_ fold changes for contrasts of interest were then calculated by subtracting the normalized log-transformed counts return by *cpm*.

#### Genome-wide screen

SgRNAs with TRIM71-specific effects were filtered, using the following contrasts:

*Contrast 1:* Unsorted cell lines (*GFP_Ctrl* vs *GFP_5xT71BS*) for differences between the genotypes that are unrelated to GFP expression:

*Contrast 2:* Sorted negative control cell lines (*GFP_Ctrl* positive vs negative) for effects on GFP expression that are unrelated to the TRIM71 pathway.

*Contrast 3:* Sorted positive control cell lines (*GFP_5xT71BS* positive vs negative) for effects on GFP expression that are both related and unrelated to the TRIM71 pathway.

*Contrast 4:* Contrast 3 - │Contrast 1│ for removing unwanted genotype effects in any direction.

*Contrast 5:* Contrast 3 – Contrast 2 for removing TRIM71-unrelated effects on GFP expression.

A z-score (‘score.up’) was calculated for contrasts 4 and 5, by subtracting the mean and dividing by the standard deviation over all sgRNAs, resulting in a ranking of sgRNAs according to their effect on GFP expression specific to the TRIM71 pathway. Effects of sgRNAs per gene were summarized by calculating the median of fold changes for all sgRNAs per gene. **Genes for the validation screen were chosen based on this z-score.**

#### Validation screen and comparison of screens

As no unsorted samples were obtained, only contrasts 2, 3, and 5 were calculated (see genome-wide screen). These contrasts were used for comparing genome-wide and validation screens.

#### Visualisation of sgRNA screens

(**Figure 1C, D**) were plotted using *Contrast 3* for negative control (‘logFC.GFP.neg.control’; x-axis) and positive control cells (‘logFC.GFP.pos.control’; y-axis) for genome-wide (**Figure 1C**) and validation (**Figure 1D**) screens. (**Figure 1E**) was plotted using *Contrast 5* (‘logFC.GFP.pathwayControlled’) for the genome-wide (x-axis) and the validation screen (y-axis). Data source: **Supplemental Table S1**.

#### RNA-seq and RIP-seq analysis

Alignment and differential gene expression analysis for RNA-seq experiments was performed as follows: Samples were aligned against the GRCm38 mouse genome assembly from GENCODE(Frankish et al., 2021), primary assembly from release M24). Alignments were performed using STAR version 2.7.3a with settings --outFilterType BySJout --outFilterMultimapNmax 20 --outMultimapperOrder Random -- alignSJoverhangMin 8 --alignSJDBoverhangMin 1 --outFilterMismatchNmax 999 -- alignIntronMin 20 --alignIntronMax 1000000 --alignMatesGapMax 1000000 -- outSAMmultNmax 1 (Gaidatzis et al., 2015). These settings follow ENCODE guidelines and report a single randomly selected alignment for reads that map to the genome no more than 20 times(Langmead et al., 2009). Gene annotations were also obtained from GENCODE release M24, transforming the gtf file into a Bioconductor TxDb object using the makeTxDbFromGFF function from the GenomicFeatures package (version 1.44.0(Lawrence et al., 2013)). Gene-level read counts were obtained by summing the counts from the union of all exons per gene, using the qCount function from the QuasR package (v1.32.0(Gaidatzis et al., 2015)) with the created TxDb and parameters orientation = ‘opposite’. EdgeR (v3.34.1(Robinson et al., 2010)) was used to test for differentially enriched genes in different conditions, using a likelihood ratio test with Benjamini-Hochberg multiple hypothesis testing correction.

#### RIP-seq normalization and identification o4f TRIM71, Ago2, and Upf1-dependent TNRC6A binding

Alignment and differential enrichment analysis for RIP-seq experiments was performed as described for RNA-seq samples(Robinson et al., 2010), with the following modifications: As we observed increased levels of most features (> 50%) in RNA IP versus input contrasts, library sizes could not be estimated using the calcNormFactors function which assumes that less than half of the features are differential. We therefore first identified features that are depleted in IP versus input (have a log_2_ fold-change less than zero) using a default normalization approach, assuming that these features can be safely assumed to correspond to non-enriched features. Next, we used the raw counts of these features (at least 3,700) for estimation of library sizes and calculation of normalization factors using calcNormFactors. The resulting normalization factors were then applied to the analysis of all features. TRIM71 and AGO2 targets were identified as transcripts with significantly reduced IP versus input TNRC6A enrichment (FDR ≤ 0.01 and log2 fold-change < 0) in *Trim71^-/-^*, or *Ago2^-/-^* compared to *wildtype* conditions, using the double contrast [(IP_Trim71^-/-^ - input_Trim71^-/-^) - (IP_wildtype – input_wildtype)], or [(IP_Ago2^-/-^ - input_Ago2^-/-^) - (IP_wildtype – input_wildtype)]. UPF1 targets were identified as transcripts with significantly reduced IP versus input TNRC6A enrichment (FDR ≤ 0.05 and log2 fold-change < 0) in *Upf1^-/-^*, compared to *wildtype* conditions, using the double contrast [(IP_Upf1^-/-^ - input_Upf1^-/-^) - (IP_wildtype – input_wildtype)].

#### Calculation of AGO2-bound hairpins, and Ago2 transcript enrichment

*AGO2 peak counts*: A file (GSE_mESC_peaks.csv.gz; GSE139345) with annotations for AGO2 peaks was downloaded from GEO(Li et al., 2020). Peaks with an adj. *P*-value of ≤ 0.05 were chosen, and peaks per gene were counted. Whole-gene AGO2 enrichments were calculated by summing all peak enrichments per gene.

#### RNA-seq heatmap

For the RNA-seq heatmap, we included transcripts with significant expression change (FDR ≤ 0.01, |log_2_(fold change)| > 1.0 in at least one condition. Expression changes were normalized to control samples (data not shown: *GFP_5xT71BS* - experiment 1; *GFP_5xT71BS* expressing *T6B_MUT_* - experiment 2).

#### Mass spectrometry analysis

Protein identification and relative quantification of the proteins was performed with MaxQuant v.1.5.3.8 using Andromeda as search engine(Cox et al., 2011), and label-free quantification (LFQ)(Cox et al., 2014; Hubner et al., 2010). The mouse subset of the UniProt v.2021_05 combined with the contaminant database from MaxQuant was searched and the protein and peptide FDR were set to 0.01.

The LFQ intensities estimated by MaxQuant were analyzed with the einprot R package (https://github.com/fmicompbio/einprot) v0.4.4. Features classified by MaxQuant as potential contaminants or reverse (decoy) hits or identified only by site, as well as features identified based on a single peptide or with a score below 10, were filtered out. The LFQ intensities were log_2_ transformed and missing values were imputed using the ’MinProb’ method from the imputeLCMD R package v2.0(Lazar, 2015) with default settings. Pairwise comparisons were performed using limma v3.50.0(Ritchie et al., 2015), considering only features with at least 2 non-imputed values across all the samples in the comparison. Estimated log_2_-fold changes and *P*-values (moderated t-test(Smyth, 2004)) from limma were used to construct volcano plots.

## Supplemental Figures

**Supplemental Figure S1.**
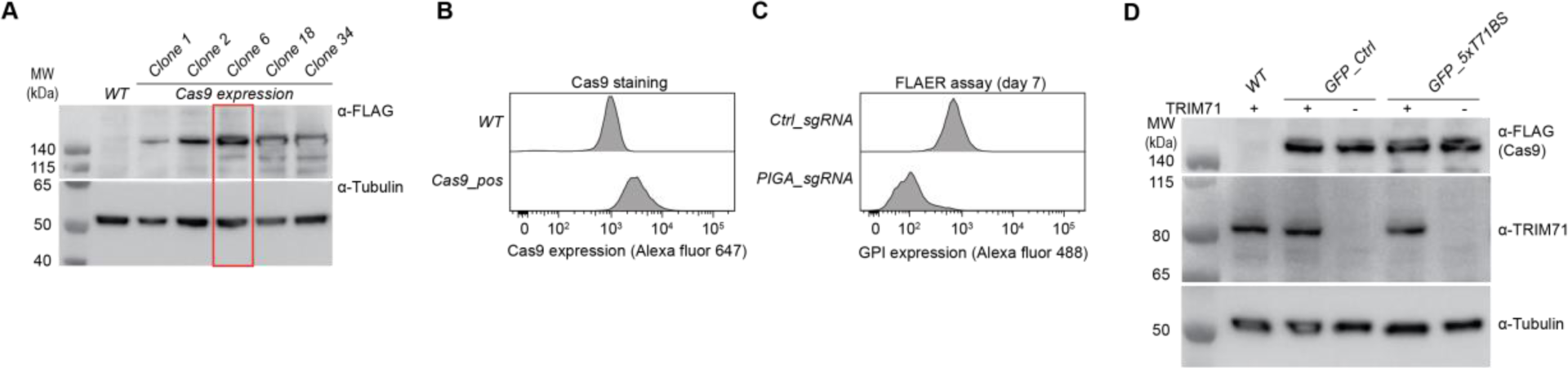
Generation of reporter cell lines for CRISPR-sgRNA-screens. **(A)** Western Blot showing Cas9 levels in different mESC clones. Clone 6 was used for further experiments (**B-D**). **(B)** Cas9 staining showing homogeneous Cas9 expression in Cas9 positive cells in comparison to *WT* cells. **(C)** FLAER assay showing high Cas9 editing efficacy in the Cas9-expressing cell line. The assay was quantified after seven days. **(D)** Western Blot showing Cas9 and TRIM71 expression levels in the generated cell lines from **Figure 1A**.

**Supplemental Figure S2.**
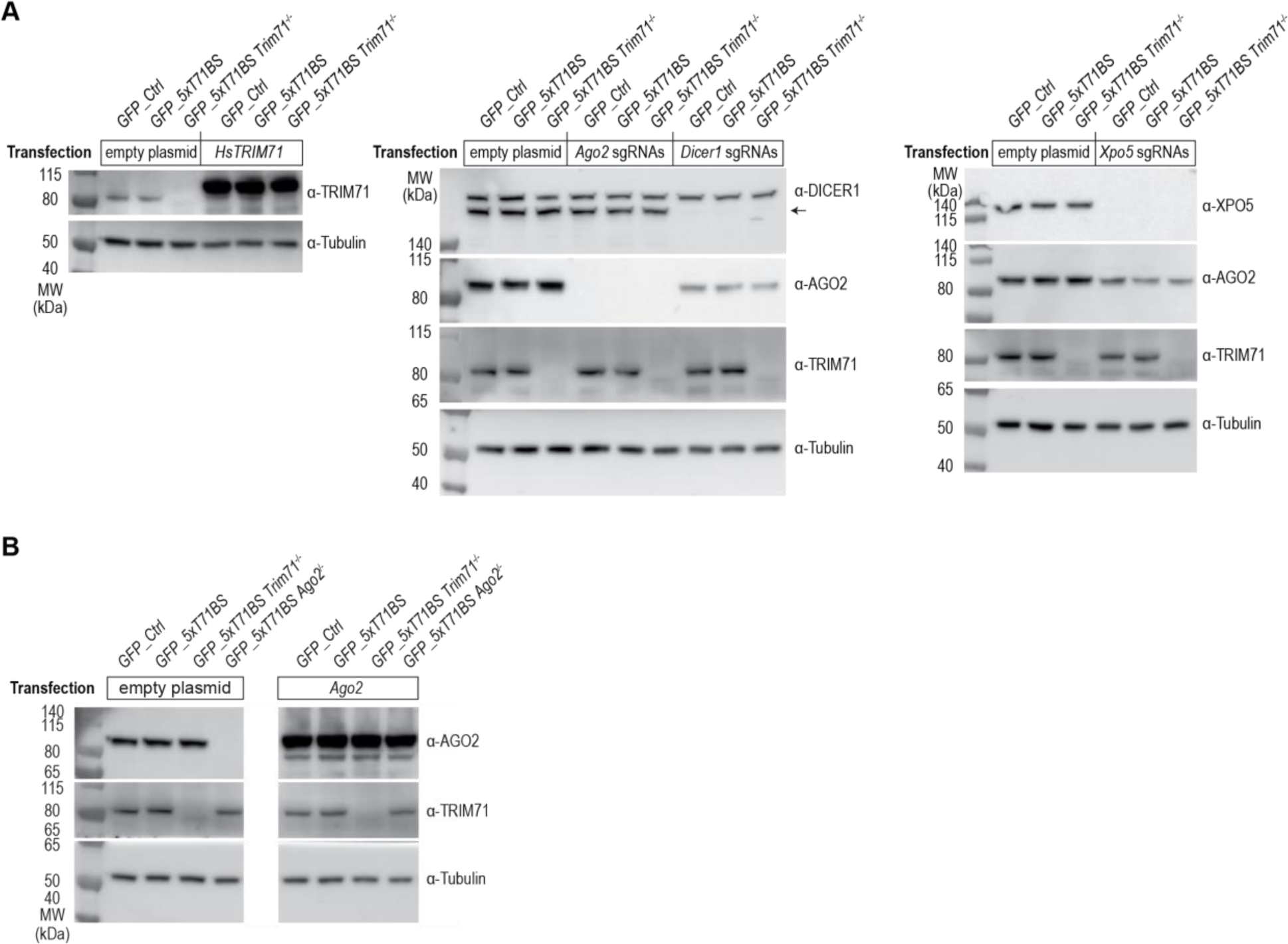
Western Blots related to Figure 2. **(A, B)** Western Blots showing the effect of transfection with *HsTRIM71* and *Ago2* overexpression plasmids, as well as sgRNAs targeting *Xpo5*, *Dicer1*, *Ago2*. Transfected cell lines are indicated. DICER1 antibodies detect two high-molecular bands. The lower one corresponding to DICER1 is marked with an arrowhead. In (**B**), white space separates lanes spliced together from the same blot (see **Supplemental Figure S8B**).

**Supplemental Figure S3.**
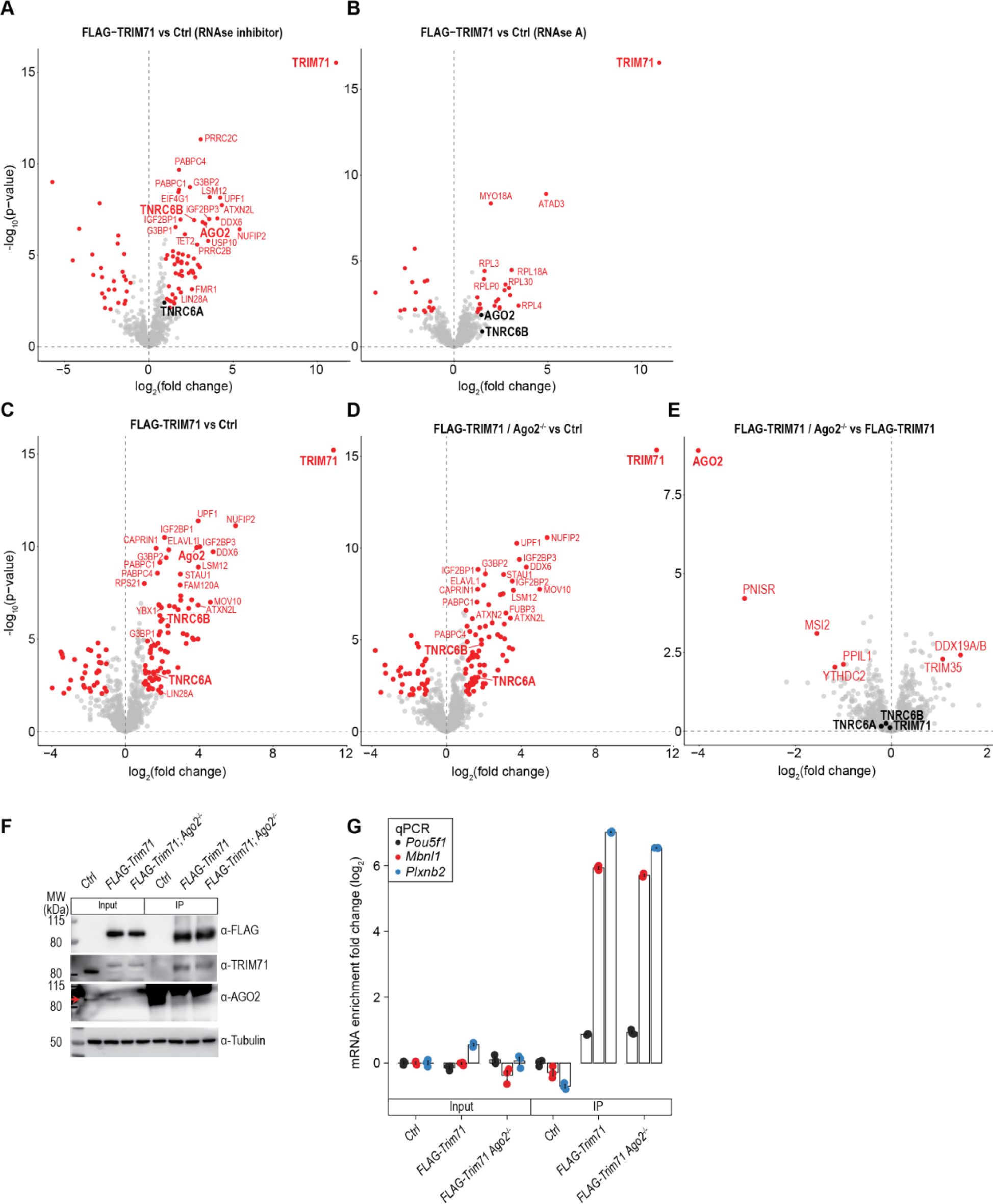
TRIM71 interactions are RNA-dependent and independent of AGO2. **(A, B)** Volcano plots showing mass spectrometry results of FLAG-AVI-TRIM71 vs untagged control cells in absence **(A)**, and presence **(B)** of RNase A (*n* = 6 biological replicates), FLAG-AVI-TRIM71 *Ago2^-/-^* cells vs FLAG-AVI-TRIM71 cells (*n* = 6 biological replicates; **[C-E]**). Significance was tested using a moderated t-test. Identified proteins with an adjusted *P* value ≤ 0.01 and |log_2_(fold change)|≥ 1 are marked in red. Proteins of interest not meeting the significance cutoff are marked in black. Data source: **Supplemental Table S2**. **(F)** Western Blot showing TRIM71, AGO2, and Tubulin expression levels for cells used **in (C-E, G).** Strong background signal from antibody chains was observed in immunoprecipitations with AGO2. AGO2 bands in input fractions are indicated with a red arrow. **(G)** Quantification of RNA immunoprecipitating with FLAG-AVI-TRIM71 by RT-qPCR in presence and absence of AGO2. Data were normalized to WT mESC input levels and represent mean ± SEM. *n* = 3 biological replicates. Background cell line (**A-G**): ‘Ctrl’; mES (129xC57BL/6) *Rosa26^Cre-ERT2/BirA-V^*.

**Supplemental Figure S4.**
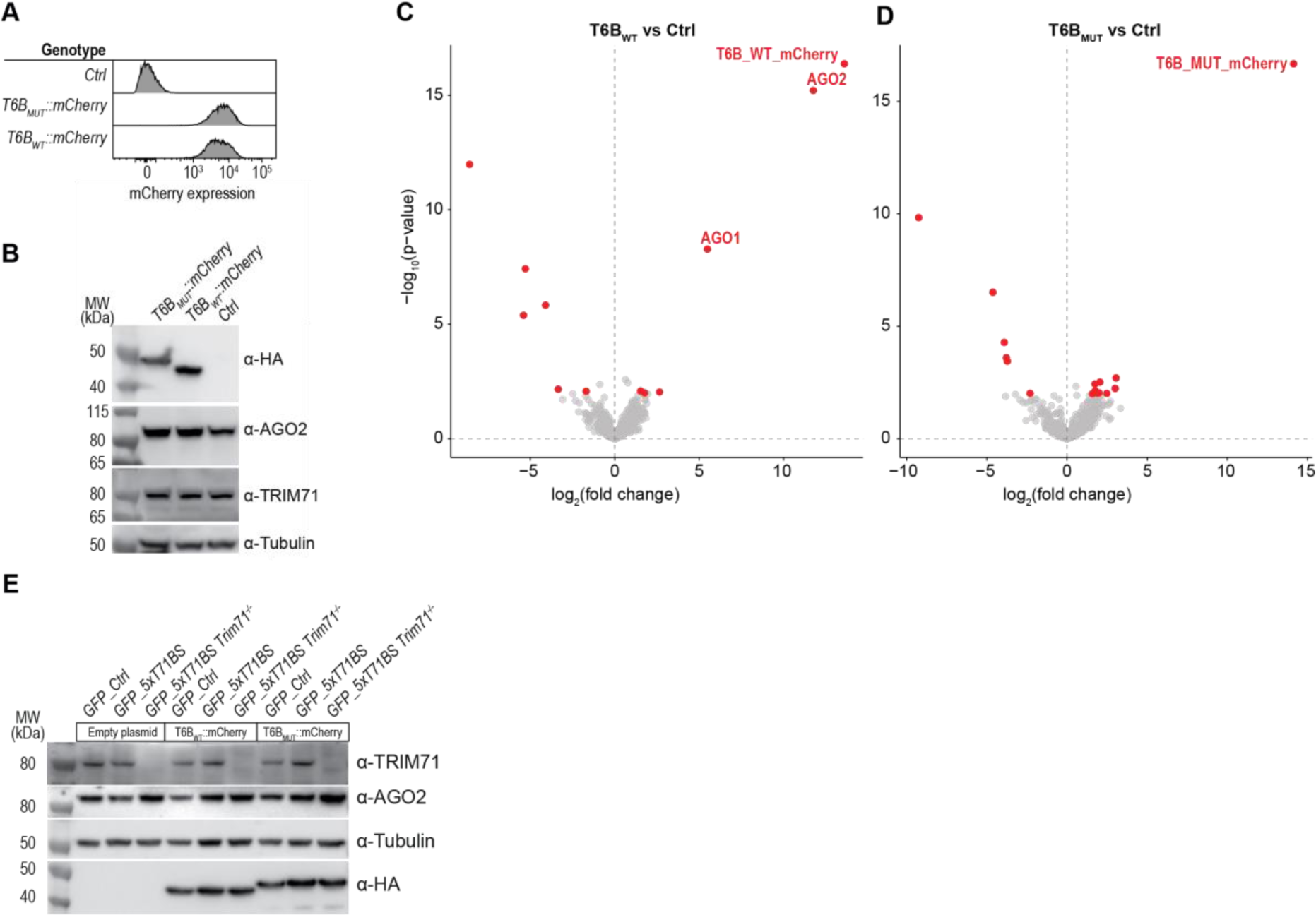
Competition of TRIM71 with AGO2 for TNRC6s. Flow cytometry plots **(A)** and Western Blot **(B)**, showing expression of untagged control cells, cells stably expressing *HA-T6B_WT_::mCherry*, and *HA-T6B_MUT_::mCherry*. **(C, D)** Volcano plots showing mass spectrometry results of HA-T6B_WT_::mCherry and HA-T6B_MUT_::mCherry immunoprecipitations (*n* = 6 biological replicates per condition). Significance was tested using a moderated t-test. Identified proteins with an adjusted *P* value ≤ 0.01 and |log_2_(fold change)|≥ 1 are marked in red. Data source: **Supplemental Table S2**. Background ‘Ctrl’ cell line (**A-D**): *GFP_5xT71BS* mESCs). **(E)** Western Blots showing overexpression of HA-T6B_WT_::mCherry and HA-T6B_MUT_::mCherry (lowest panel), and corresponding TRIM71, AGO2, and Tubulin levels of cells used in (**Figure 4F**).

**Supplemental Figure S5.**
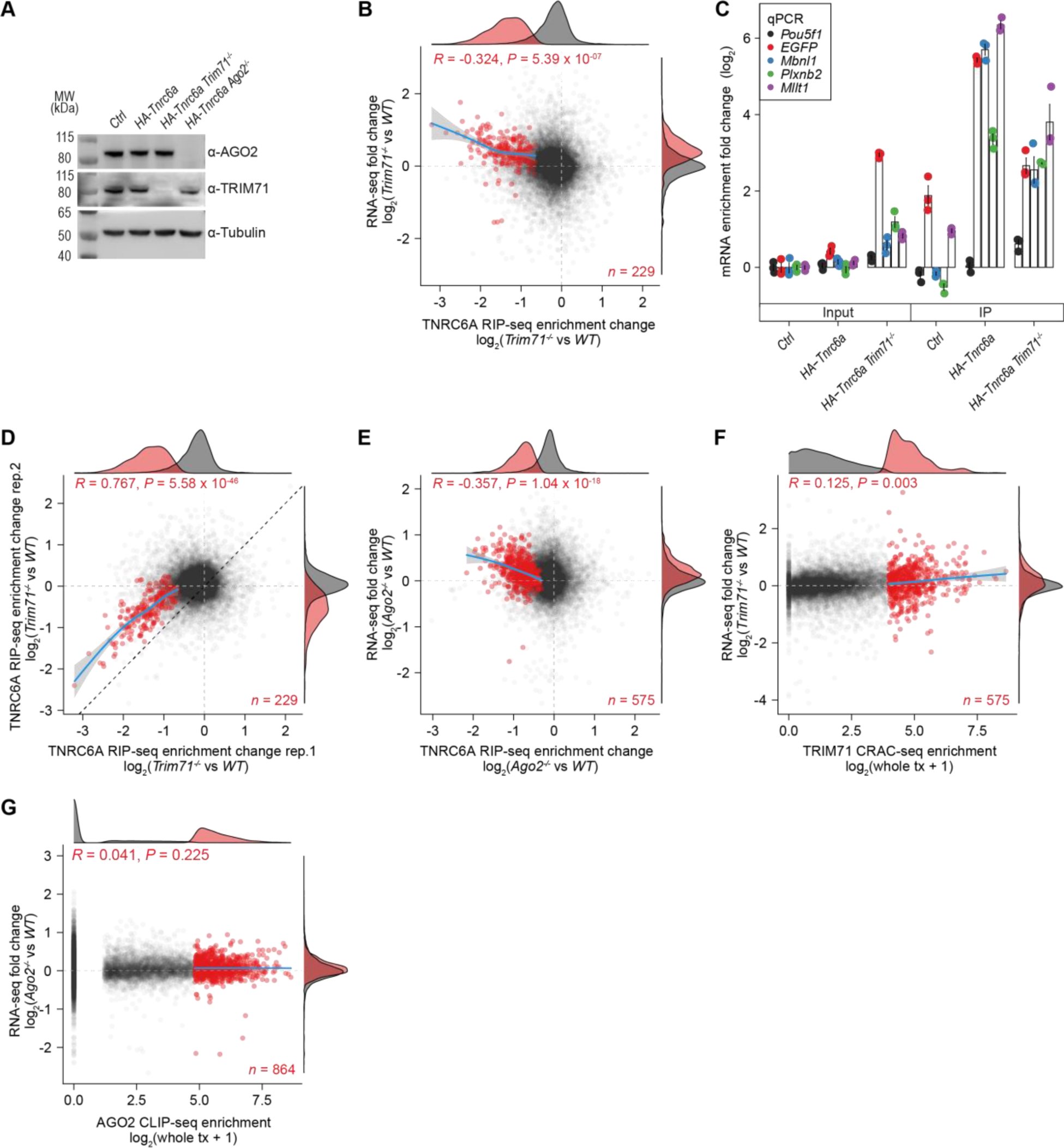
TNRC6A binds to TRIM71 target transcripts in the presence of TRIM71. **(A)** Western Blot showing TRIM71, AGO2, and Tubulin expression levels in cell lines used for RNA-immunoprecipitation experiments. **(B)** Scatter plot showing changes of TNRC6A transcript enrichment in *Trim71^-/-^* vs *WT* cells as quantified by RIP-seq (x-axis), and changes in transcript level in *Trim71^-/-^* vs *WT* cells as quantified by RNA-seq (y axis). Red color indicates transcripts with significant loss of RIP-seq enrichment (FDR ≤ 0.01, log_2_[fold change] < 0). A generalized additive model was fitted to these transcripts (blue line, 95% confidence interval: grey shadow). *R* indicates Pearson’s correlation coefficient calculated for transcripts with significant loss of enrichment. **(C)** Bar plot showing results of an RNA-immunoprecipitation experiment using untagged control cells, (‘*Ctrl*’), *HA-AVI-Tnrc6a* cells, *HA-AVI-Tnrc6a* cells lacking *Trim71* (*HA-Tnrc6a Trim71^-/-^*). Data were normalized to input *Ctrl* cells and represent mean ± SEM. *n* = 3 biological replicates. **(D)** Scatter plot showing changes of TNRC6A transcript enrichment in *Trim71^-/-^* vs *WT* cells as quantified by RIP-seq in rep. 1 (x-axis), and rep. 2 (y-axis). Red color indicates transcripts with significant TNRC6A enrichment change in rep. 1. *R* indicates Pearson’s correlation coefficient for transcripts with significant TNRC6A enrichment change in rep. 1. **(E)** Scatter plot showing changes of TNRC6A transcript enrichment in *Ago2^-/-^* vs *WT* cells as quantified by RIP-seq (x-axis), and changes in transcript level in *Ago2^-/-^* vs *WT* cells as quantified by RNA-seq (y axis). Red color indicates transcripts with significant loss of RIP-seq enrichment in *Ago2^-/-^* vs *WT* cells (FDR ≤ 0.01, log_2_[fold change] < 0). A generalized additive model was fitted to these transcripts (blue line, 95% confidence interval: grey shadow). *R* is Pearson’s correlation coefficient calculated for transcripts with significant loss of enrichment. The corresponding density distributions for (**B, D, E**) are shown on top (x-axis), and to the right (y-axis). *P* values in (**B, D, E**) are calculated using a paired, two-sided t test for correlation. Background cell line (**B-E**): *GFP_5xT71BS* mESCs. (**F**) Scatter plot showing TRIM71 transcript binding assessed by CRAC-seq (x-axis: log_2_[enrichment + 1]; (Welte et al., 2019)) and changes in transcript levels in *Trim71^-/-^* vs *WT* cells as quantified by RNA-seq (y axis). Red color indicates the 5% most strongly bound transcripts. (**G**) Scatter plot showing AGO2 transcript binding assessed by CLIP-seq (x-axis: log_2_[enrichment + 1]; (Li et al., 2020)) and changes in transcript levels in *Ago2^-/-^* vs *WT* cells as quantified by RNA-seq (y axis). Red color indicates the 5% most strongly bound transcripts. Data source (**A-G**): **Supplemental Table S3**.

**Supplemental Figure S6.**
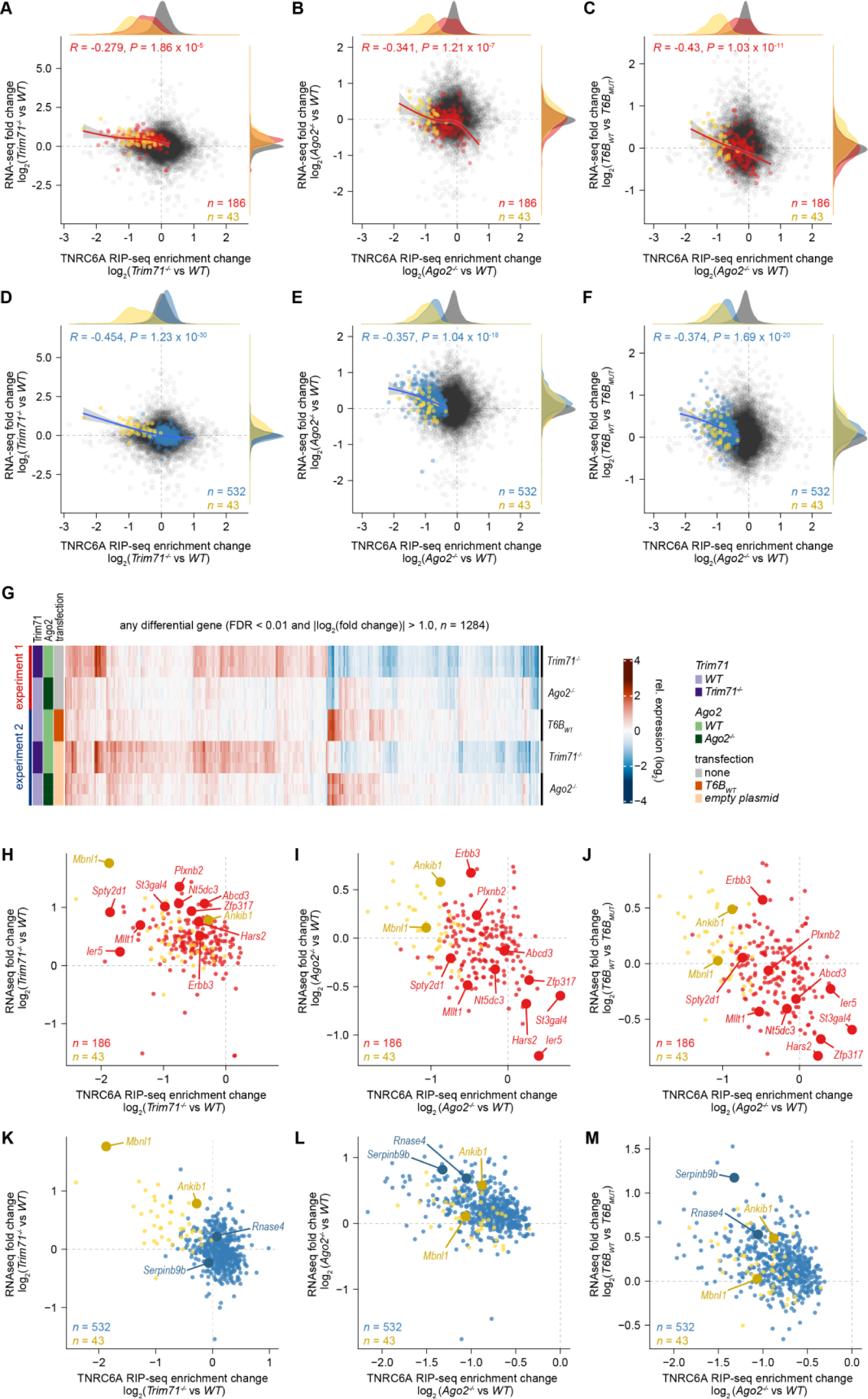
TNRC6A RIP-seq identifies TRIM71 and AGO2 target transcripts with high precision. Exclusive TRIM71 target transcripts are silenced more efficiently in *Ago2* knock-out cells. **(A, D)** Scatter plots showing changes of TNRC6A transcript enrichment in *Trim71^-/-^* vs *WT* cells as quantified by RIP-seq (x-axis), and changes in transcript level in *Trim71^-/-^* vs *WT* cells as quantified by RNA-seq (y axis). **(B, E)** Scatter plots showing changes of TNRC6A transcript enrichment in *Ago2^-/-^* vs *WT* cells as quantified by RIP-seq (x-axis), and changes in transcript level in *Ago2^-/-^* vs *WT* cells as quantified by RNA-seq (y axis). **(C, F)** Scatter plots showing changes of TNRC6A transcript enrichment in *Ago2^-/-^* vs *WT* cells as quantified by RIP-seq (x-axis), and changes in transcript level in *T6B_WT_* vs *T6B_MUT_* cells as quantified by RNA-seq (y axis). <u>Red color</u> indicates transcripts with significant loss of RIP-seq enrichment in *Trim71^-/-^* vs *WT* cells (FDR ≤ 0.01, log_2_[fold change] < 0), but not in *Ago2^-/-^* vs *WT* cells. <u>Blue color</u> indicates transcripts with significant loss of RIP-seq enrichment in *Ago2^-/-^*vs *WT* cells (FDR ≤ 0.01, log_2_[fold change) < 0], but not in *Trim71^-/-^* vs *WT* cells. <u>Yellow color</u> indicates transcripts with significant loss of RIP-seq enrichment in *Trim71^-/-^* vs *WT* cells, <u>and</u> in *Ago2^-/-^* vs *WT* cells. **(G**) Heatmap showing gene expression fold changes of differentially expressed genes (FDR ≤ 0.01, |log_2_(fold change)| > 1 in *Trim71^-/-^*, *Ago2^-/-^* (experiment 1), and *T6B_WT_*, *Trim71^-/-^*, and *Ago2^-/-^* cells (experiment 2). Cells in experiment 2 were transfected with either *T6B_WT_*, *T6B_MUT_* or empty plasmid (for *Trim71^-/-^*, and *Ago2^-/-^* cells), and selected with 1 µg/ml Puromycin for two days, followed by RNA extraction and sequencing. Individual replicates are normalized to the mean of three *WT* cells (for experiment 1), and T6B_MUT_ expressing cells (experiment 2), respectively. **(H-M)** Scatter plots showing transcripts chosen for luciferase assay validation experiments. Coloring as in **(A-F)**. *R* values are Pearson’s correlation coefficients, calculated for red and yellow (**A-C**), or blue and yellow (**D-F**) data points. *P* values are calculated using a paired, two-sided t test for correlation. Data source **(A-F, H-M)**: **Supplemental Table S3**. Background cell line: *GFP_5xT71BS* mESCs.

**Supplemental Figure S7.**
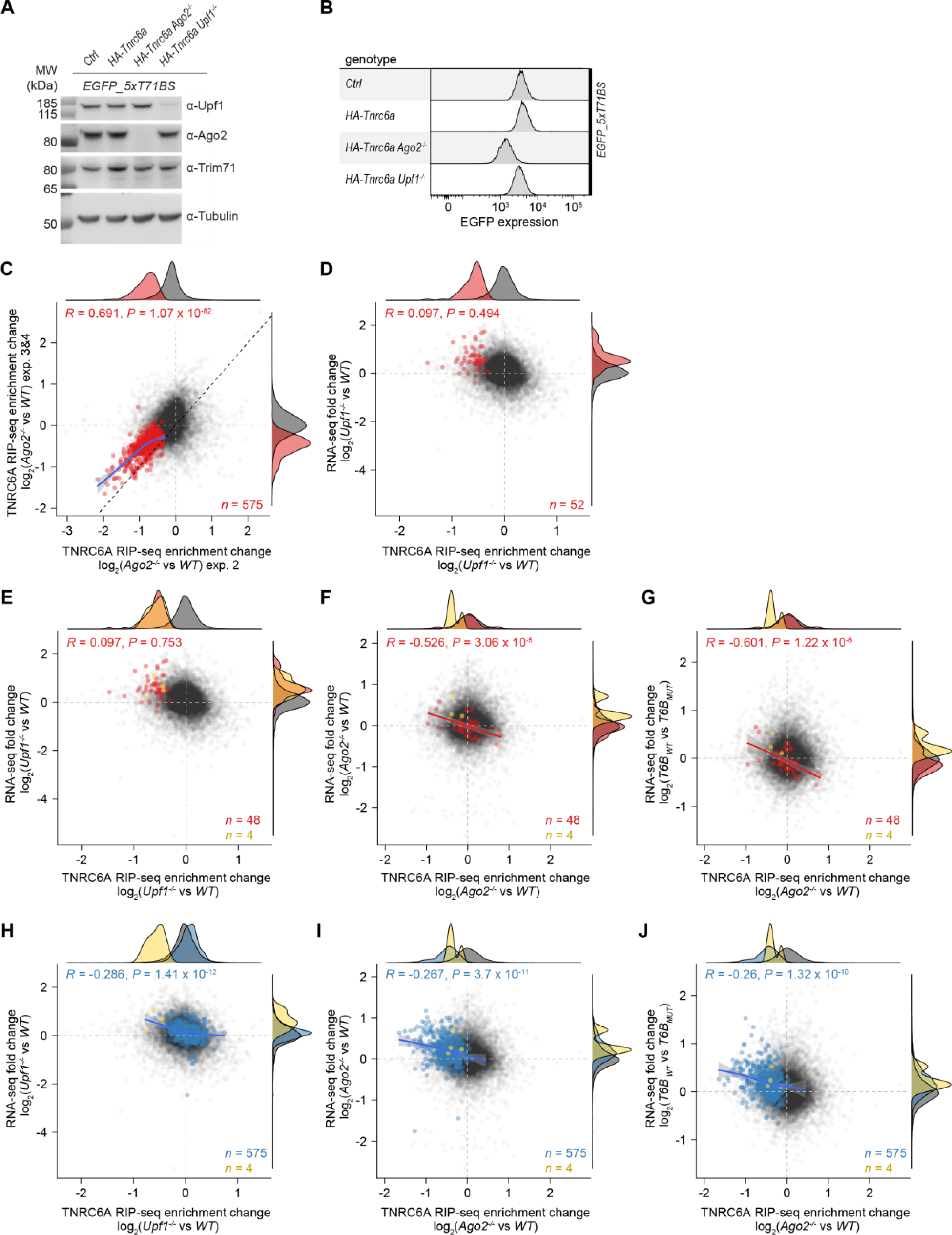
TNRC6A RIP-seq identifies UPF1 target transcripts. Exclusive UPF1 target transcripts are silenced more efficiently in *Ago2* knock-out cells. **(A)** Western Blot showing UPF1, AGO2, and Tubulin expression levels in cell lines used for RIP-seq experiments. **(B)** Flow cytometry analysis, showing expression of GFP_5xT71BS in cells used for RIP-seq. **(C)** Scatter plot showing changes of TNRC6A transcript enrichment in *Ago2^-/-^* vs *WT* cells as quantified by RIP-seq in exp. 2 (x-axis), and exp. 3&4 (y-axis). Red color indicates transcripts with significant TNRC6A enrichment change in *Ago2^-/-^* vs *WT* cells (FDR ≤ 0.01, log_2_[fold change] < 0) in exp. 2. *R* indicates Pearson’s correlation coefficient for transcripts with significant loss of TNRC6A enrichment in exp. 2. **(D)** Scatter plot showing changes of TNRC6A transcript enrichment in *Upf1^-/-^* vs *WT* cells as quantified by RIP-seq (x-axis), and changes in transcript level in *Upf1^-/-^* vs *WT* cells as quantified by RNA-seq (y axis). Red color indicates transcripts with significant loss of RIP-seq enrichment (FDR ≤ 0.05, log_2_[fold change] < 0. *R* indicates Pearson’s correlation coefficient calculated for transcripts with significant loss of enrichment. **(E, H)** Scatter plots showing changes of TNRC6A transcript enrichment in *Upf1^-/-^* vs *WT* cells as quantified by RIP-seq (x-axis), and changes in transcript level in *Upf1^-/-^* vs *WT* cells as quantified by RNA-seq (y axis). **(F, I)** Scatter plots showing changes of TNRC6A transcript enrichment in *Ago2^-/-^* vs *WT* cells as quantified by RIP-seq (x-axis), and changes in transcript level in *Ago2^-/-^* vs *WT* cells as quantified by RNA-seq (y axis). **(G, J)** Scatter plots showing changes of TNRC6A transcript enrichment in *Ago2^-/-^* vs *WT* cells as quantified by RIP-seq (x-axis), and changes in transcript level in *T6B_WT_* vs *T6B_MUT_* cells as quantified by RNA-seq (y axis). <u>Red color</u> indicates transcripts with significant loss of TNRC6A RIP-seq enrichment in *Upf1^-/-^* vs *WT* cells (FDR ≤ 0.05, log_2_[fold change] < 0), but not in *Ago2^-/-^* vs *WT* cells. <u>Blue color</u> indicates transcripts with significant loss of RIP-seq enrichment in *Ago2^-/-^* vs *WT* cells (FDR ≤ 0.01, log_2_[fold change) < 0], but not in *Upf1^-/-^* vs *WT* cells. <u>Yellow color</u> indicates transcripts with significant loss of RIP-seq enrichment in *Upf1^-/-^* vs *WT* cells, <u>and</u> in *Ago2^-/-^* vs *WT* cells. R values are Pearson’s correlation coefficients. *P* values are calculated using a paired, two-sided t test for correlation. Data source **(C-J)**: **Supplemental Table S3**. Background cell line: *GFP_5xT71BS* mESCs.

## Supplemental uncropped gels

### Black boxed areas are shown in corresponding figures

**Supplemental uncropped gels Figure 1.**
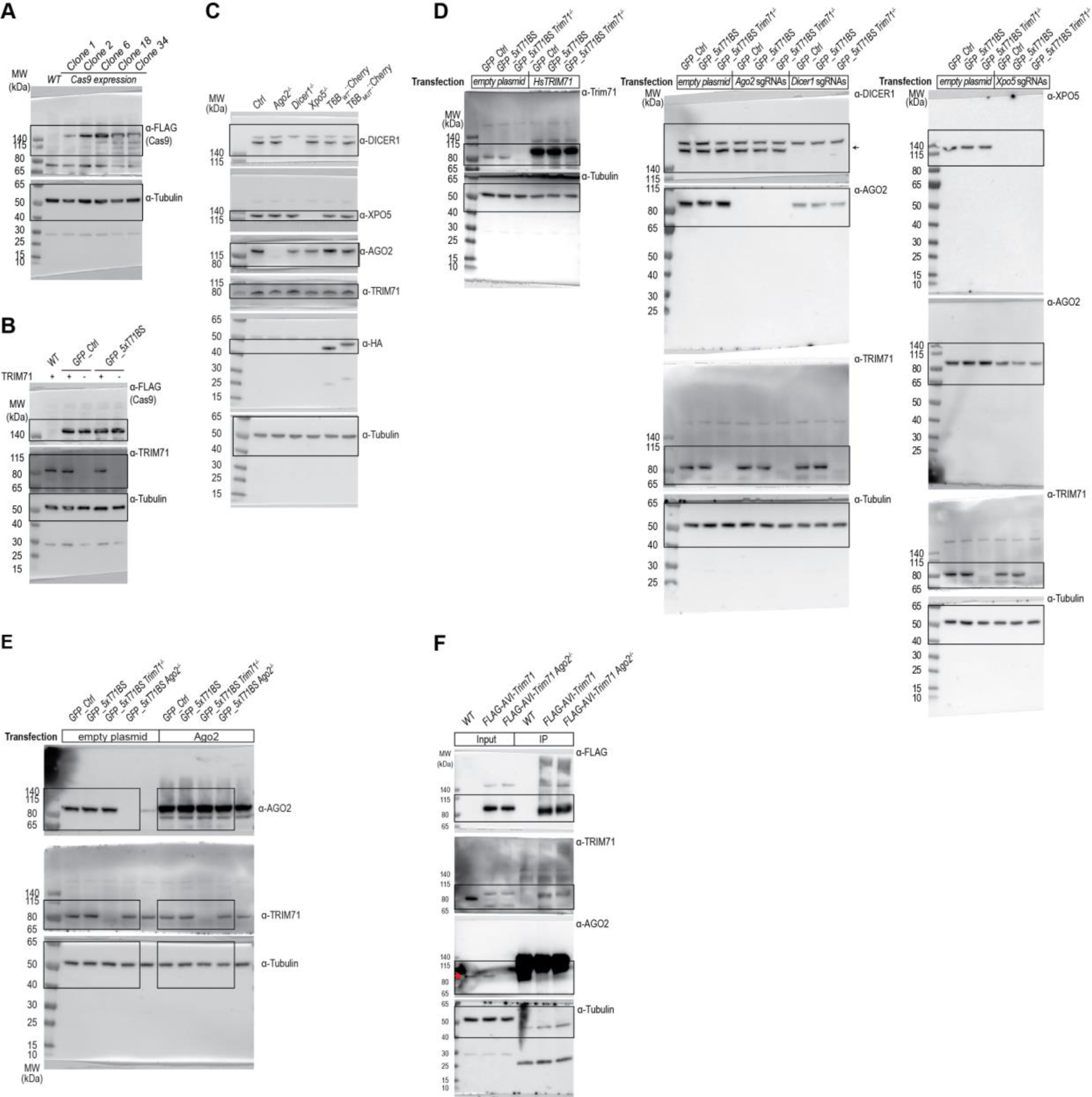
**(A)** Gels from **Supplemental Figure S1A**. **(B)** Gels from **Supplemental Figure S1D**. **(C)** Gels from **Figure 2A**. **(D)** Gels from **Supplemental Figure S2A**. **(E)** Gels from **Supplemental Figure S2B**. **(F)** Gels from **Supplemental Figure S3F**.

**Supplemental uncropped gels Figure 2.**
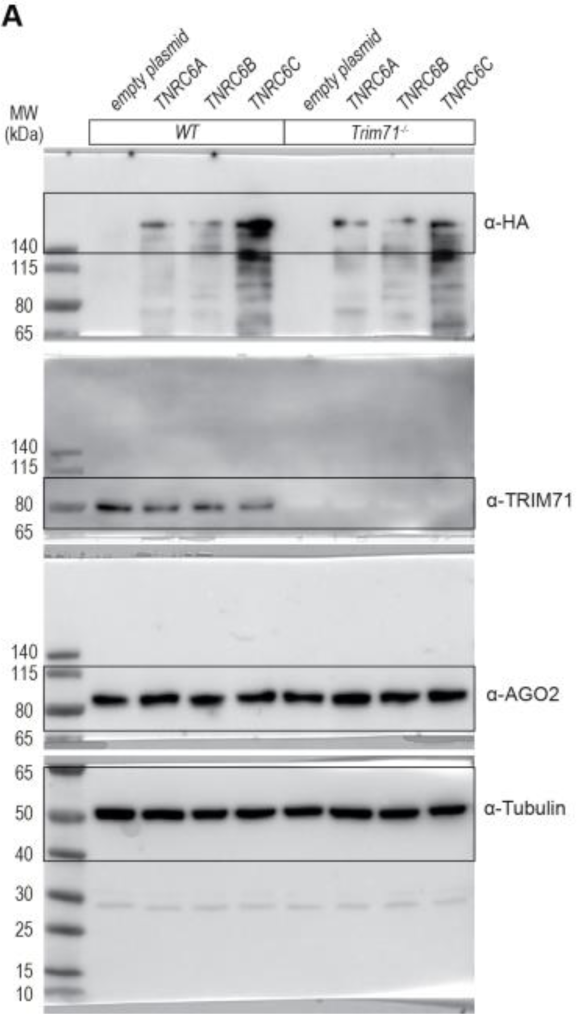
**(A)** Gels from **Figure 3B**.

**Supplemental uncropped gels Figure 3.**
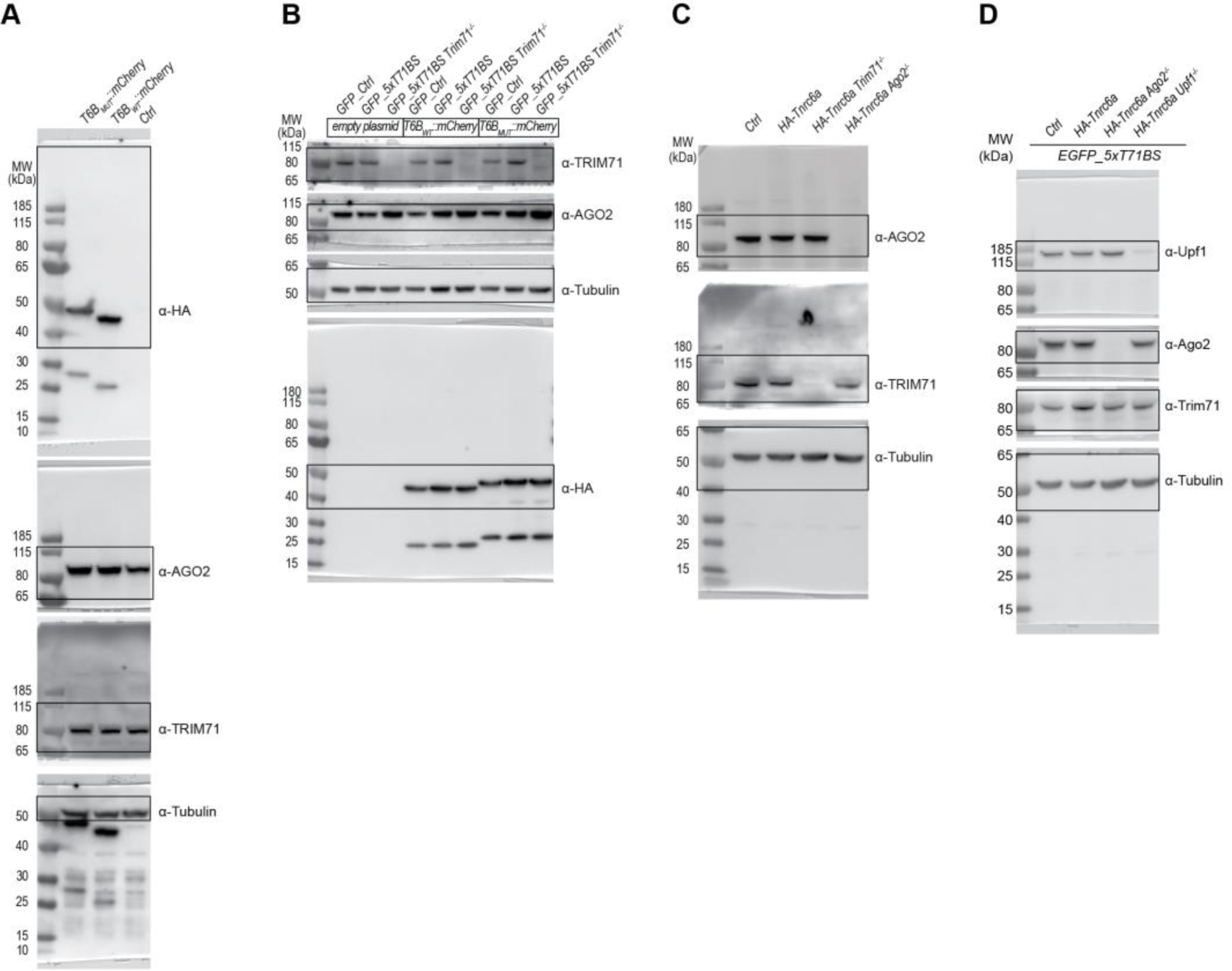
**(A)** Gels from **Supplemental Figure S4B**. **(B)** Gels from **Supplemental Figure S4E**. **(C)** Gels from **Supplemental Figure S5A**. **(D)** Gels from **Supplemental Figure S7A**.

## Extended Data Tables

**Supplemental Table S1. SgRNA screen results. Related to Figure 1B-D.**

<u>Tab 1</u>: genome-wide screen results by sgRNA. <u>Tab 2</u>: genome-wide screen results by gene. <u>Tab 3</u>: validation screen results by sgRNA. <u>Tab 4</u>: validation screen results by gene. <u>Tab 5</u>: comparison of genome-wide and validation screens. Results by gene. <u>Tab 6</u>: SgRNA library used for the validation screen. <u>Legend Tab 1-6</u>: logFC = log_2_ fold change between groups; logFC.genotype.unspecific: *GFP_5xT71BS* day 8 unsorted vs *GFP_Ctrl* day 8 unsorted. logFC.GFP.neg.control: *GFP_Ctrl* day 8 (GFP high) – *GFP_Ctrl* day 8 (GFP low). logFC.GFP.pos.control: *GFP_5xT71BS* day 8 (GFP high) – *GFP_5xT71BS* day 8 (GFP low). logFC.GFP.genotypeControlled: logFC.GFP.pos.control - │logFC.genotype.unspecific│. logFC.GFP.pathwayControlled: logFC.GFP.pos.control - logFC.GFP.neg.control. score.up: z-score of logFC.GFP.genotypeControlled and logFC.GFP.pathwayControlled. score.down: negative z-score of logFC.GFP.genotypeControlled and logFC.GFP.pathwayControlled. p.value = two-sided *P*-value; FDR = Benjamini Hochberg adjusted *P*-value). <u>Tab 7, 8</u>: Raw read counts for each sgRNA in genome-wide (Tab 7), and validation (Tab 8) sgRNA screens. <u>Legend Tab 7, 8</u>: column ‘sample_type’: mESCs = count data generated from sequencing screened mESCs, plasmid = count data generated from sequencing input sgRNA library; column ‘genotype’: genotype of mESCs; column ‘sorting_status’: sorted_GFP_high = 25% high GFP FACS-sorted mESCs, sorted_GFP_low = 25% low GFP FACS-sorted mESCs, unsorted = unsorted mESCs; column ‘timepoint’: day_8 = DNA extraction at day 8; day_4 = DNA extraction at day 4.

**Supplemental Table S2. Mass spectrometry results. Related to Figure 3a-c and Supplemental Figure S4A-E, Supplemental Figure S4C, D.**

<u>Legend</u>: Protein.IDs: UniProt protein IDs, Protein.names: Protein names, Gene.names: Gene names, logFC: Log2 fold change, AveExpr: Average expression, P.Value: two-sided P-value (moderated t-test), adj.P.Val: Benjamini Hochberg adjusted P-value, mlog10p: −log_10_(adj.P.val), iBAQ.avg: average IBAQ value, iBAQ.sd: standard deviation IBAQ value.

**Supplemental Table S3. RNA-seq and RIP-seq results. Related to Figure 5–7, Supplemental Figure S5-7.**

<u>Legend:</u> LFC, Log_2_ fold change; FDR, Benjamini Hochberg adjusted *P*-value. <u>Blue columns:</u> Differential gene expression analysis with EdgeR of *Trim71^-/-^* vs *WT* mESCs (Trim71_KO_vs_WT), *Ago2^-/-^* vs *WT* mESCs (AGO2_KO_vs_WT), *Upf1^-/-^* vs *WT* mESCs (UPF1_KO_vs_WT), and *T6B_WT_* vs *T6B_MUT_* mESCs (T6B_WT_vs_MUT; *n* = 3 biological replicates). <u>Yellow column:</u> Trim71.CRAC.enrichment.vs.RNA: Whole transcript CRAC enrichment. **Data source:** (Welte et al., 2019). <u>Orange column:</u> Ago2_CLIP_peaks: Ago2 CLIP peaks per gene. Ago2_CLIP_enrichment: Sum of Ago2 CLIP peaks per gene. **Data source:** (Li et al., 2020). <u>Green columns</u>: RIP-seq results: *n* = 4 independent experiments, 3 replicates per experiment: RIP_1, RIP_2, RIP_3&4. LFC_IP-enrichment HA-Tnrc6a: log_2_(IP vs input [Tnrc6a_IP_input]). Dependent_Tnrc6a_IP_input: TRUE, if FDR_Tnrc6a_IP_input ≤ 0.01. LFC_Tnrc6a_RIP_enr.change_Trim71_KO_vs_WT: log_2_(enrichment difference of HA-Tnrc6a IP vs input in Trim71^-/-^ vs WT mESCs). Dependent_enr.change_Trim71_KO_vs_WT: TRUE, if FDR_Tnrc6a_RIP_enr.change_Trim71_KO_vs_WT ≤ 0.01. LFC_Tnrc6a_RIP_enr.change_Ago2_KO_vs_WT: log_2_(enrichment difference of HA-Tnrc6a IP vs input in Ago2^-/-^ vs WT mESCs). Dependent_enr.change_Ago2_vs_WT: TRUE, if FDR_Tnrc6a_RIP_enr.change_Ago2_KO_vs_WT ≤ 0.01. LFC_Tnrc6a_RIP_enr.change_Upf1_KO_vs_WT: log_2_(enrichment difference of HA-Tnrc6a IP vs input in Upf1^-/-^ vs WT mESCs). Dependent_enr.change_Upf1_KO_vs_WT: TRUE, if FDR_Tnrc6a_RIP_enr.change_Upf1_KO_vs_WT ≤ 0.05. target_to_pathway_TRIM71_AGO2: none, RIP_1_dependent_enr.change_Trim71_KO_vs_WT *and* RIP_2_dependent_enr.change_Ago2_KO_vs_WT FALSE. Trim71-only: RIP_1_dependent_enr.change_Trim71_KO_vs_WT: TRUE *and* RIP_2_dependent_enr.change_Ago2_KO_vs_WT: FALSE. Ago2-only: RIP_2_dependent_enr.change_Ago2_KO_vs_WT: TRUE *and* RIP_1_dependent_enr.change_Trim71_KO_vs_WT: FALSE. Common: RIP_1_dependent_enr.change_Trim71_KO_vs_WT *and* RIP_2_dependent_enr.change_Ago2_KO_vs_WT TRUE. target_to_pathway_UPF1_AGO2: none, RIP_dependent_enr.change_Upf1_KO_vs_WT *and* RIP_2_dependent_enr.change_Ago2_KO_vs_WT FALSE. Upf1-only: RIP_3&4_dependent_enr.change_Upf1_KO_vs_WT: TRUE *and* RIP_2_dependent_enr.change_Ago2_KO_vs_WT: FALSE. Ago2-only: RIP_2_dependent_enr.change_Ago2_KO_vs_WT: TRUE *and* RIP_3&4_dependent_enr.change_Upf1_KO_vs_WT: FALSE. Common: RIP_3&4_dependent_enr.change_Upf1_KO_vs_WT *and* RIP_2_dependent_enr.change_Ago2_KO_vs_WT TRUE

**Supplemental Table S4.**
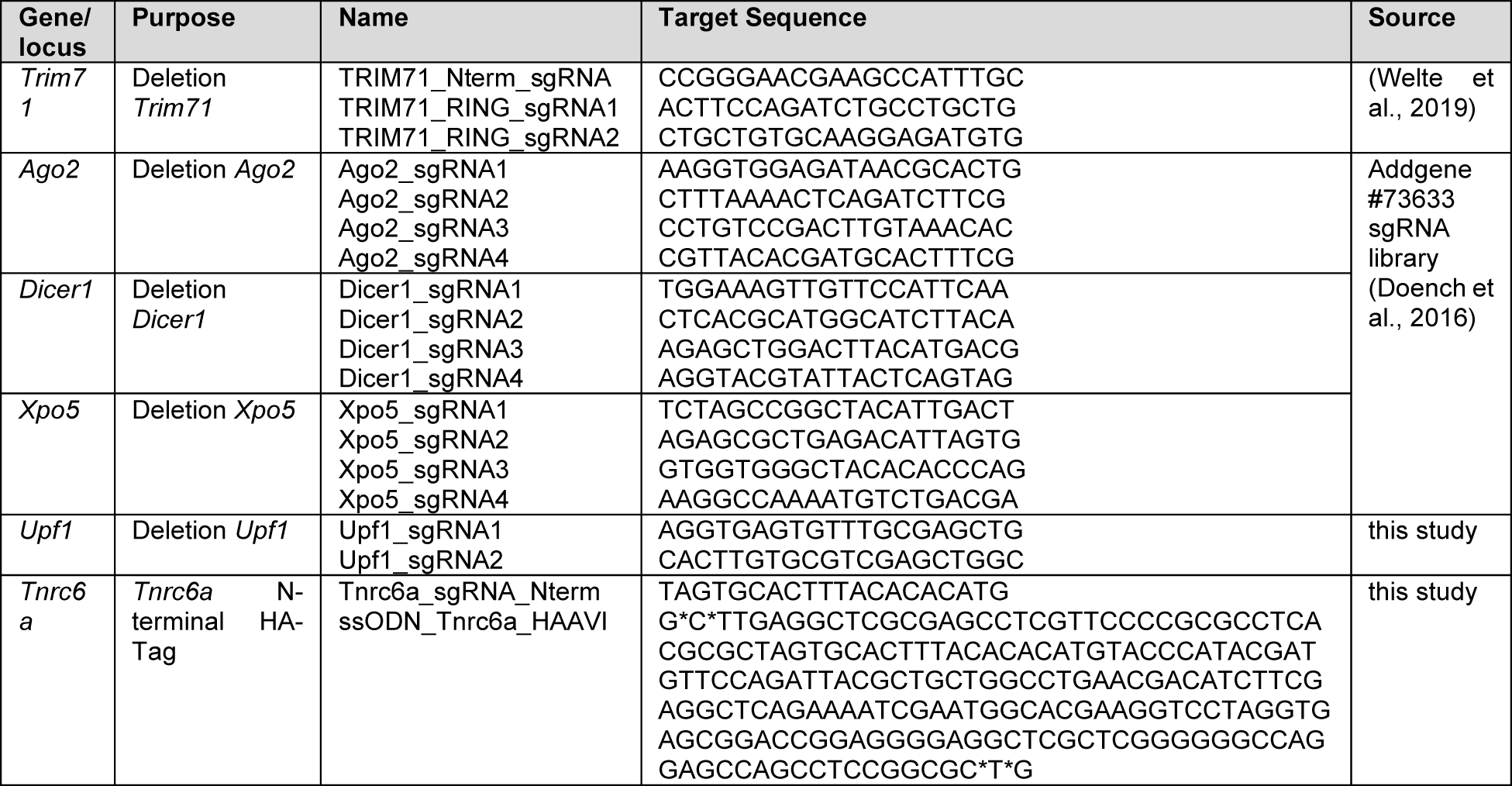
sgRNA sequences and ssODNs. Asterisks in ssODN sequences indicate phosporothioate bonds.

**Supplemental Table S5.** Cell lines.

| Cell line No. | Genotype | Description | Source |
| --- | --- | --- | --- |
| 1 | mES (129xC57BL/6) *Rosa26*<sup>Cre-ERT2/BirA-V5</sup> | Untagged control cell line for Trim71-immunoprecipitations | (Ostapczuk et al., 2018) |
| 4 | mES (129xC57BL/6) *Rosa26*<sup>Cre-ERT2/BirA-V5</sup> *Trim71*<sup>FlagAvi/FlagAvi</sup> | N terminal 3xFLAG-AVI-tagged *Trim71* | (Welte et al., 2019) |
| 20 | mES (V6.5), *Ago2*<sup>Halo-LSL/+</sup> | N-terminal FLAG-tagged *Ago2* | (Li et al., 2020) |
| 21 | mES (129S6/SvEvTac) | Parental cell line for Cas9 integration | (Lienert et al., 2011) |
| 22 | mES (129S6/SvEvTac), stable transfection *Cas9::T2A::Blasticidin* | Overexpression of Cas9, parental cell line for GFP reporter gene integration | this study |
| 23 | mES (129S6/SvEvTac), *yGlob* (*GFP_Ctrl*), stable transfection *Cas9::T2A::Blasticidin* | Overexpression of Cas9, overexpression of *GFP_Ctrl* | this study |
| 24 | mES (129S6/SvEvTac), *yGlob* (*GFP_5xT71BS*), stable transfection *Cas9::T2A::Blasticidin* | Overexpression of Cas9, overexpression of *GFP_5xT71BS* | this study |
| 25 | mES (129S6/SvEvTac), *yGlob* (*GFP_Ctrl*), *Trim71*<sup>-/-</sup>, stable transfection *Cas9::T2A::Blasticidin* | Overexpression of Cas9, overexpression of *GFP_Ctrl*, deletion of *Trim71* | this study |
| 26 | mES (129S6/SvEvTac), *yGlob* (*GFP_5xT71BS*), *Trim71*<sup>-/-</sup>, stable transfection *Cas9::T2A::Blasticidin* | Overexpression of Cas9, overexpression of *GFP_5xT71BS*, deletion of *Trim71* | this study |
| 27 | mES (129S6/SvEvTac), *yGlob* (*GFP_5xT71BS*), *Ago2*<sup>-/-</sup>, stable transfection *Cas9::T2A::Blasticidin* | Overexpression of Cas9, overexpression of *GFP_5xT71BS*, Deletion of *Ago2* | this study |
| 28 | mES (129S6/SvEvTac), *yGlob* (*GFP_5xT71BS*), *Dicer1*<sup>-/-</sup>, stable transfection *Cas9::T2A::Blasticidin* | Overexpression of Cas9, overexpression of *GFP_5xT71BS*, deletion of *Dicer1* | this study |
| 29 | mES (129S6/SvEvTac), *yGlob* (*GFP_5xT71BS*), *Xpo5*<sup>-/-</sup>, | Overexpression of Cas9, overexpression of *GFP_5xT71BS*, | this study |
|  | stable transfection *Cas9::T2A::Blasticidin* | deletion of *Xpo5* |  |
| 34 | mES (129S6/SvEvTac), *yGlob (GFP_5xT71BS)*, *HA-AVI-Tnrc6a*, stable transfection *Cas9::T2A::Blasticidin* | Overexpression of *Cas9*, overexpression of *GFP_5xT71BS*, N-terminal HA-AVI-tag of *Tnrc6a* | this study |
| 35 | mES (129S6/SvEvTac), *yGlob (GFP_5xT71BS)*, *HA-AVI-Tnrc6a*, *Trim71<sup>-/-</sup>*, stable transfection *Cas9::T2A::Blasticidin* | Overexpression of *Cas9*, overexpression of *GFP_5xT71BS*, N-terminal HA-AVI-tag of *Tnrc6a*, deletion of *Trim71* | this study |
| 36 | mES (129S6/SvEvTac), *yGlob (GFP_5xT71BS)*, *HA-AVI-Tnrc6a*, *Ago2<sup>-/-</sup>*, stable transfection *Cas9::T2A::Blasticidin* | Overexpression of *Cas9*, overexpression of *GFP_5xT71BS*, N-terminal HA-AVI-tag of *Tnrc6a*, deletion of *Ago2* | this study |
| 37 | mES (129S6/SvEvTac), *yGlob (GFP_5xT71BS)*, stable transfection *FH-T6B<sub>WT</sub>::mCherry*, stable transfection *Cas9::T2A::Blasticidin* | Overexpression of *Cas9*, overexpression of *GFP_5xT71BS*, Overexpression of *FH-T6B<sub>WT</sub>::mCherry* | this study |
| 38 | mES (129S6/SvEvTac), *yGlob (GFP_5xT71BS)*, stable transfection *FH-T6B<sub>MUT</sub>::mCherry*, stable transfection *Cas9::T2A::Blasticidin* | Overexpression of *Cas9*, overexpression of *GFP_5xT71BS*, Overexpression of *FH-T6B<sub>MUT</sub>::mCherry* | this study |
| 39 | mES (129S6/SvEvTac), *yGlob (GFP_5xT71BS)*, *HA-AVI-Tnrc6a*, *Upf1<sup>-/-</sup>*, stable transfection *Cas9::T2A::Blasticidin* | Overexpression of *Cas9*, overexpression of *GFP_5xT71BS*, N-terminal HA-AVI-tag of *Tnrc6a*, deletion of *Upf1* | this study |

**Supplemental Table S6A.** Overexpression plasmids.

| Name | Backbone | Resistance | Insert | Backbone reference | Insert reference | Primers used |
| --- | --- | --- | --- | --- | --- | --- |
| pTW71 | pC2P | AMP | *Cas9::BSD* | pTW42 (Welte et al., 2019) | pEF1-B; Matyas Flemr, Marc Bühler | 225-228 |
| pTW97 | pL1_1L | AMP | *GFP_Ctrl (EGFP::unc-54 3'UTR)* | pL1_1L; (Lienert et al., 2011) | pTW47; (Welte et al., 2019) | 406, 407 |
| pTW98 | pL1_1L | AMP | *GFP_5xT71BS (EGFP::mab-10<sub>condensed</sub>)* | pL1_1L; (Lienert et al., 2011) | pTW48; (Welte et al., 2019) | 406, 407 |
| pTW110 | pCIP | AMP | *mmuAgo2 IRES2 Puro* | Addgene #79009 (Young et al., 2016) | Addgene #73538 (Ngondo et al., 2018) | 745, 746 |
| pTW111 | pCIP | AMP | *HsTNRC6A IRES2 Puro* | Addgene #79009 (Young et al., 2016) | Addgene #19883 (Landthaler et al., 2008) | 747, 748 |
| pTW112 | pCIP | AMP | *HsTNRC6B IRES2 Puro* | Addgene #79009 (Young et al., 2016) | Addgene #10979 (Meister et al., 2005) | 747, 750, 780, 781 |
| pTW113 | pCIP | AMP | *HsTNRC6C IRES2 Puro* | Addgene #79009 (Young et al., 2016) | Addgene #19885 (Landthaler et al., 2008) | 751, 752 |
| pTW114 | pCIP | AMP | *FH-T6B<sub>WT</sub>::mCherry IRES2 Puro* | Addgene #79009 | pIRESneo FLAG/HA-T6B_WT- | 764, 765, 768, 769 |
|  |  |  |  | (Young et al., 2016) | eYFP; (Hauptmann et al., 2015); pTW51; (Welte et al., 2019) |  |
| pTW115 | pCiP | AMP | *FH_T6B<sub>MUT</sub>::mCherry IRES2 Puro* | Addgene #79009 (Young et al., 2016) | pIRESneo FLAG/HA-T6B_MUT-eYFP; (Hauptmann et al., 2015); pTW51; (Welte et al., 2019) | 764, 765, 768, 769 |

**Supplemental Table S6B.**
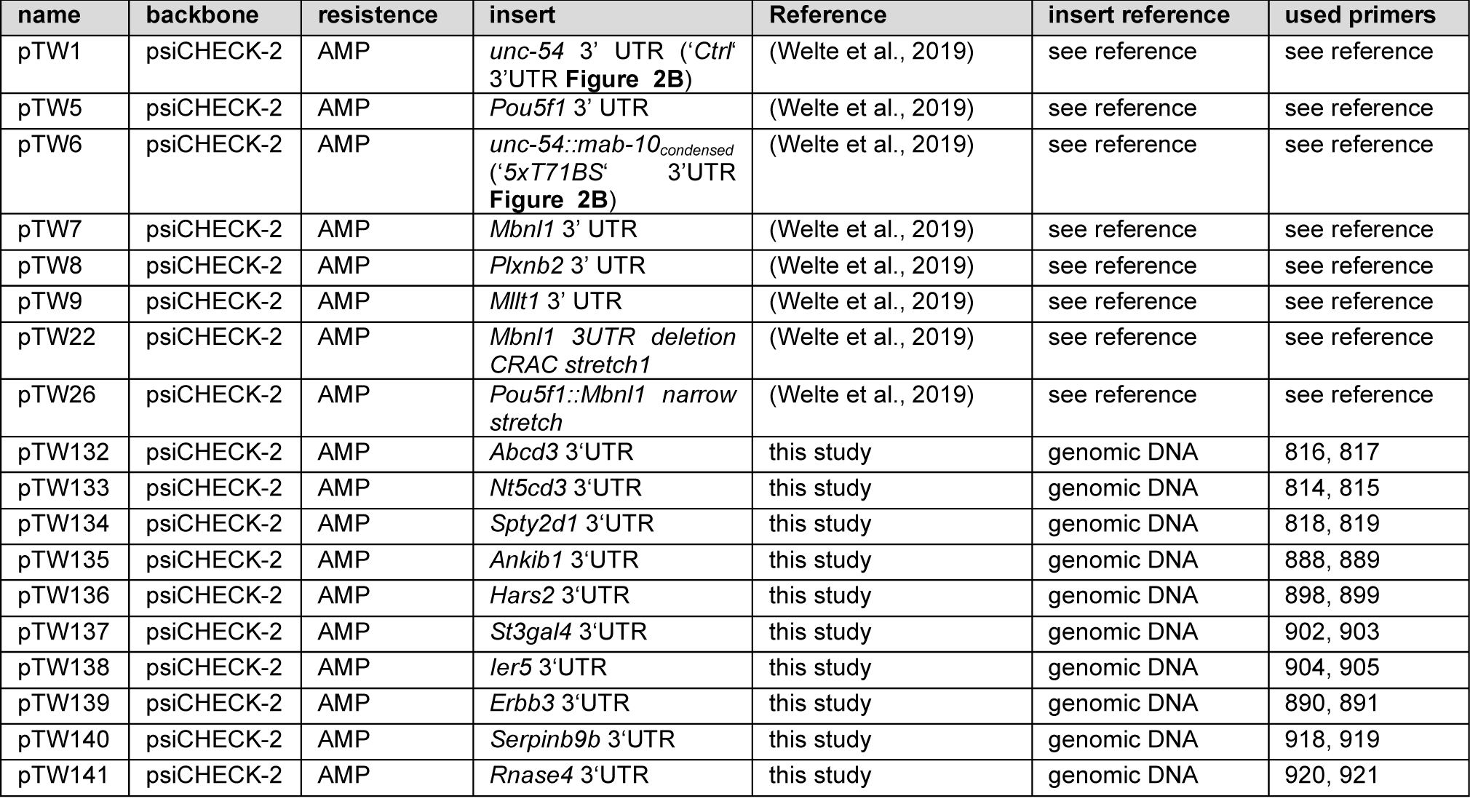
Luciferase reporter gene constructs.

**Supplemental Table S7A.**
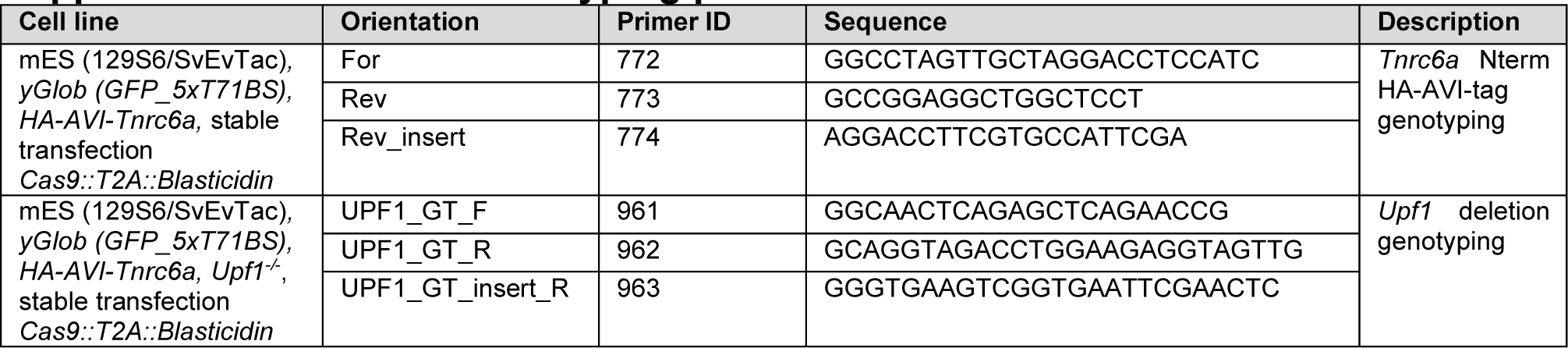
Genotyping primers.

**Supplemental Table S7B.**
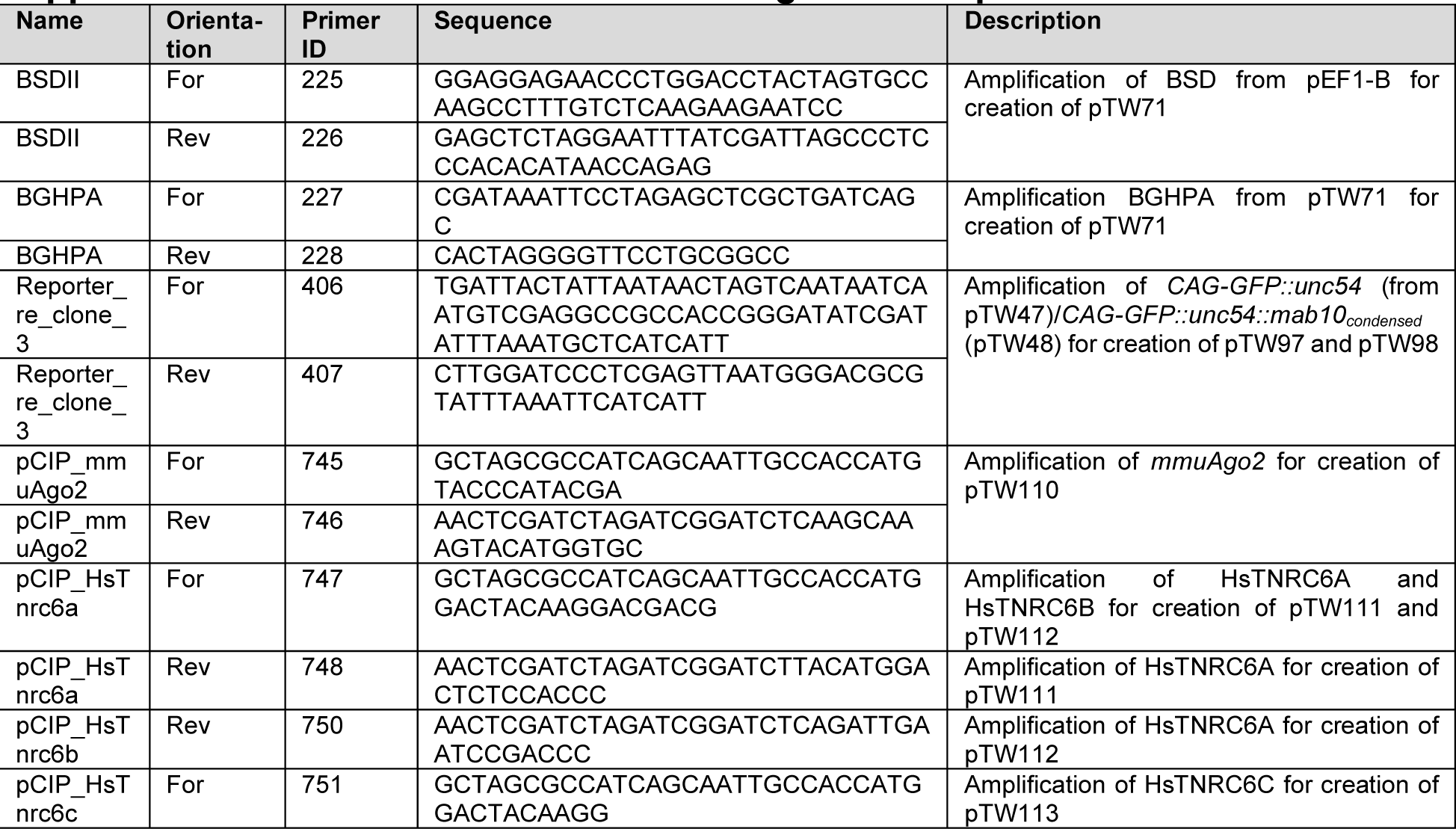
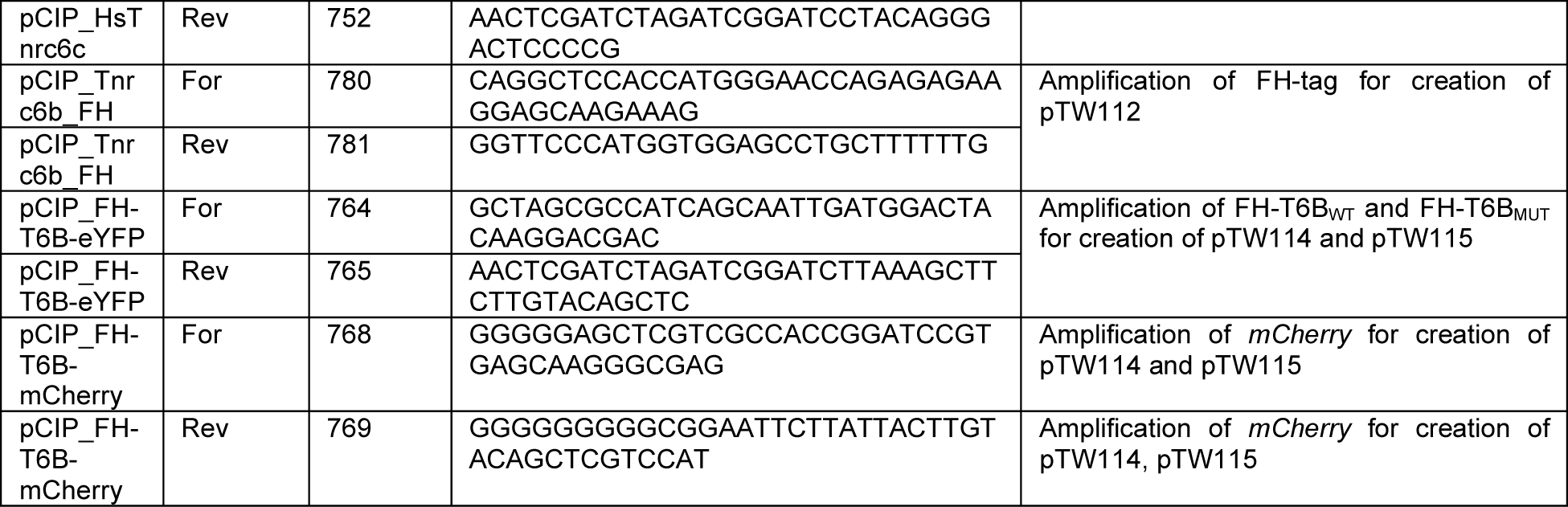
Primers for cloning of overexpression constructs.

**Supplemental Table S7C.**
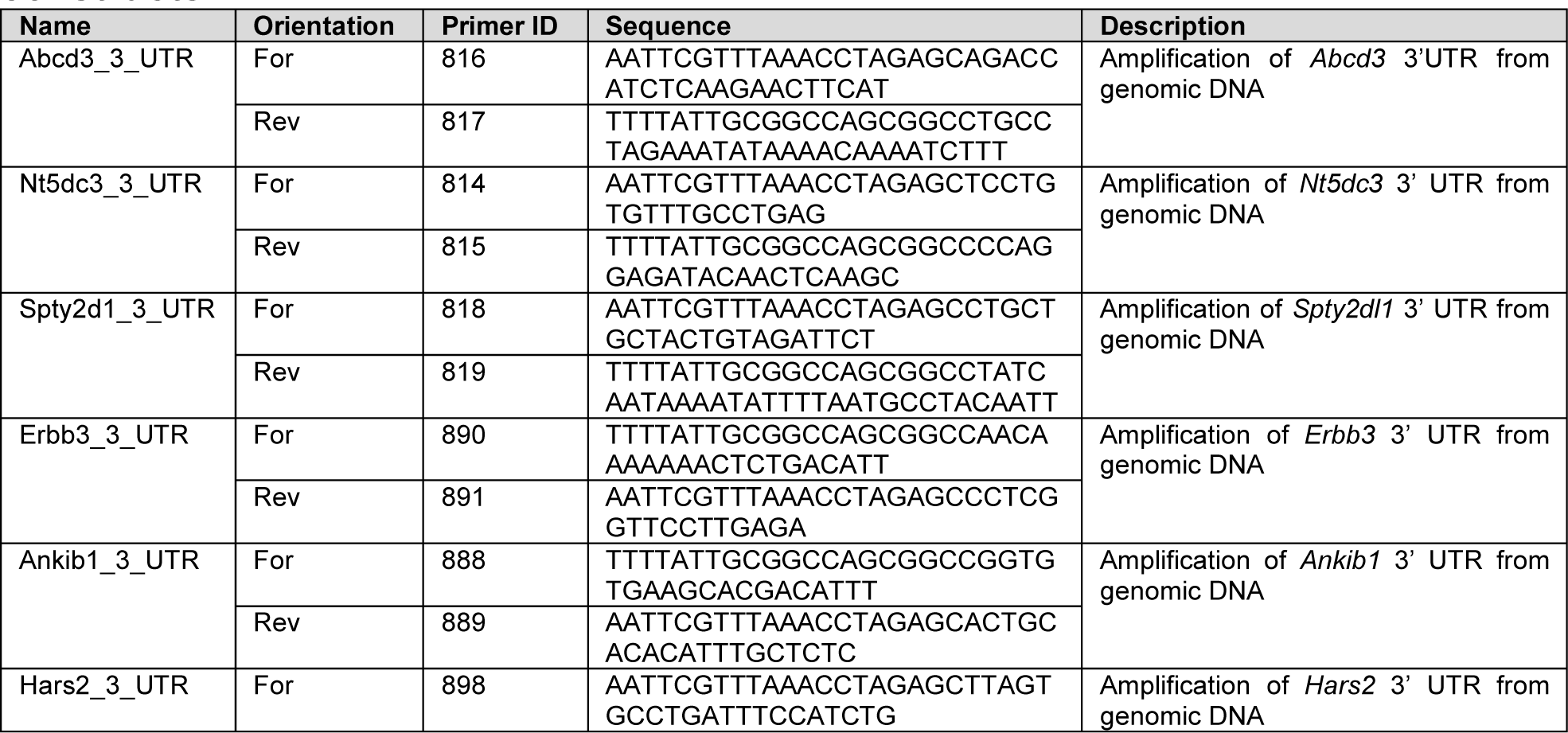
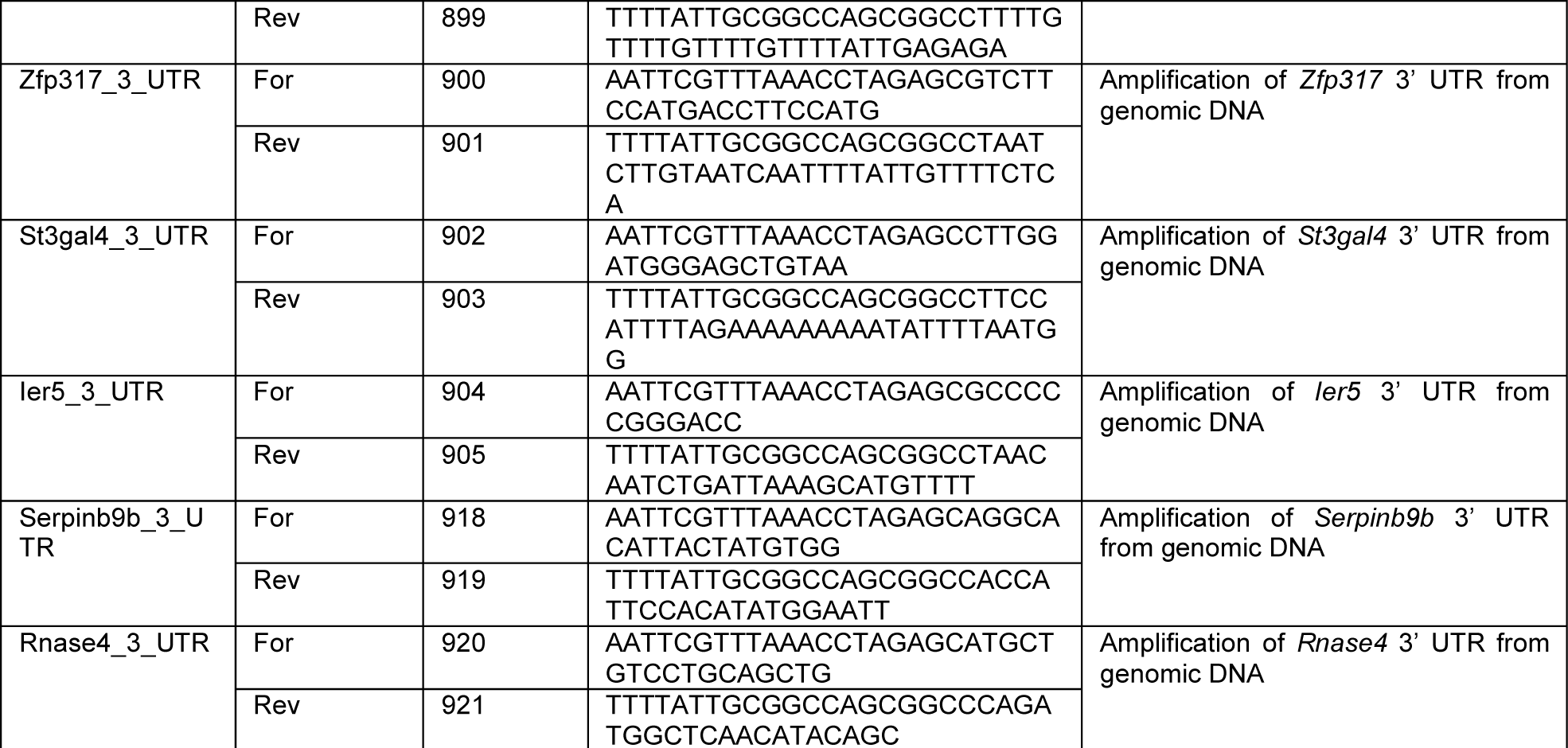
Primers for cloning of luciferase reporter gene constructs.

**Supplemental Table S8.**
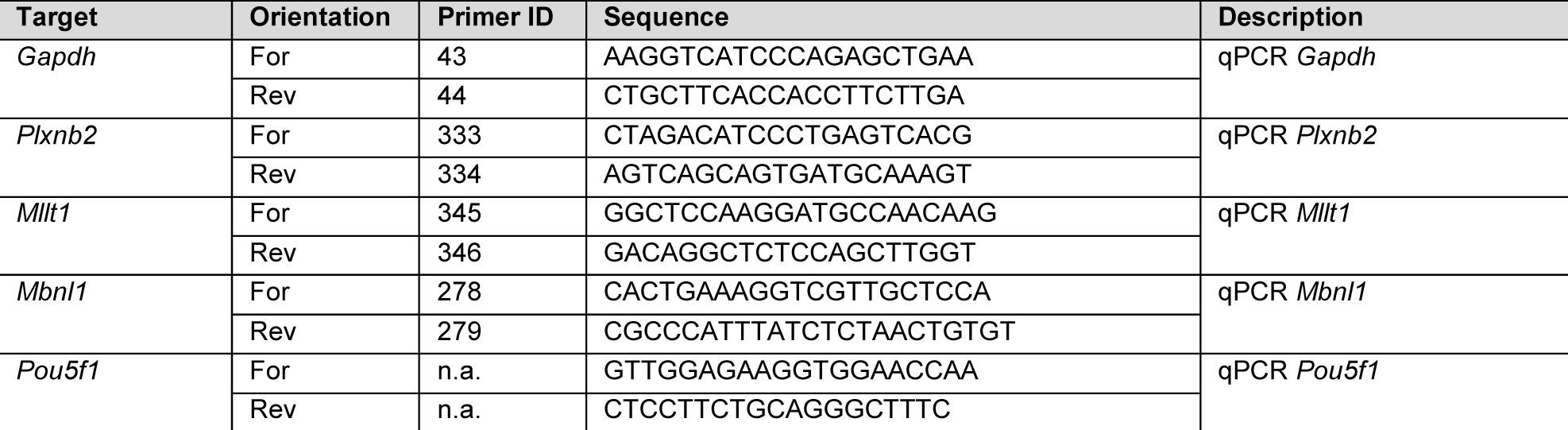
qPCR Primers.

**Supplemental Table S9.** Antibodies for Western Blotting.

| Antibody | Dilution | Incubation | Company | Catalog No. |
| --- | --- | --- | --- | --- |
| α-TRIM71 | 1:500 | 4 °C overnight | R&D Systems | AF5104 |
| α-Sheep-HRP | 1:5,000 | 1h room temperature | Abcam | ab97130 |
| α-Tubulin | 1:10,000 | 1h room temperature | Abcam | ab6160 |
| α-Rat-HRP | 1:15,000 | 1h room temperature | Jackson ImmunoResearch | 112035175 |
| α-AGO2 | 1:1,000 | 1h room temperature | Abcam | ab156870 |
| α-DICER1 | 1:1,000 | 1h room temperature | Abcam | ab227518 |
| α-XPO5 | 1:1,000 | 1h room temperature | Abcam | ab168844 |
| α-HA | 1:500 | 1h room temperature | MerckMillipore | 11867423001 |
| α-Rabbit-HRP | 1:5,000 | 1h room temperature | GE Healthcare | LNA934V |
| α-FLAG-HRP | 1:500 | 1h room temperature | Sigma | A8592 |
| α-UPF1 | 1:2,000 | 1h room temperature | Abcam | ab86057 |

